# Trade-offs between cost and information in cellular prediction

**DOI:** 10.1101/2023.01.10.523390

**Authors:** Age J. Tjalma, Vahe Galstyan, Jeroen Goedhart, Lotte Slim, Nils B. Becker, Pieter Rein ten Wolde

## Abstract

Living cells can leverage correlations in environmental fluctuations to predict the future environment and mount a response ahead of time. To this end, cells need to encode the past signal into the output of the intracellular network from which the future input is predicted. Yet, storing information is costly while not all features of the past signal are equally informative on the future input signal. Here, we show, for two classes of input signals, that cellular networks can reach the fundamental bound on the predictive information as set by the information extracted from the past signal: pushpull networks can reach this information bound for Markovian signals, while networks that take a temporal derivative can reach the bound for predicting the future derivative of non-Markovian signals. However, the bits of past information that are most informative about the future signal are also prohibitively costly. As a result, the optimal system that maximizes the predictive information for a given resource cost is, in general, not at the information bound. Applying our theory to the chemotaxis network of *Escherichia coli* reveals that its adaptive kernel is optimal for predicting future concentration changes over a broad range of background concentrations, and that the system has been tailored to predicting these changes in shallow gradients.

---

Single-celled organisms live in a highly dynamic environment to which they continually have to respond and adapt. To this end, they employ a range of response strategies, tailored to the temporal structure of the environmental variations. When these variations are highly regular, such as the daily light variations, it becomes beneficial to develop a clock from which the time and hence the current and future environment can be inferred [1, 2]. In the other limit, when the fluctuations are entirely unpredictable, cells have no choice but to resort to either the strategy of detect-and-respond or the bet-hedging strategy of stochastic switching between different phenotypes [3]. Yet arguably the most fascinating strategy lies in between these two extremes. When the environmental fluctuations happen with some regularity, then it becomes feasible to predict the future environment and initiate a response ahead of time. While it is commonly believed that only higher organisms can predict the future, experiments have vividly demonstrated that even single-cell organisms can leverage temporal correlations in environmental fluctuations in order to predict, e.g., future nutrient levels [4, 5].

The ability to predict future signals can provide a fitness benefit [6]. The capacity to anticipate changes in oxygen levels [4], or the arrival of sugars or stress signals [5], can increase the growth rate of single-celled organisms; modeling has revealed that prediction can enhance bacterial chemotaxis [7]. Yet, a predict-and-anticipate strategy is only advantageous if the cell can reliably predict the future on timescales that are longer than the time it takes to mount a response. What fundamentally limits the accuracy of cellular prediction remains, however, poorly understood.

While the cell needs to predict the future environment, it can only sense the present and remember the past (Fig. 1A). Consequently, for a given amount of information the cell can store about the present and past signal, there is a maximum amount of information it can possibly have about the future [6, 8] (Fig. 1C-I). This *information bound* is determined by the temporal structure of the environmental fluctuations [8, 9].

**FIG. 1.**
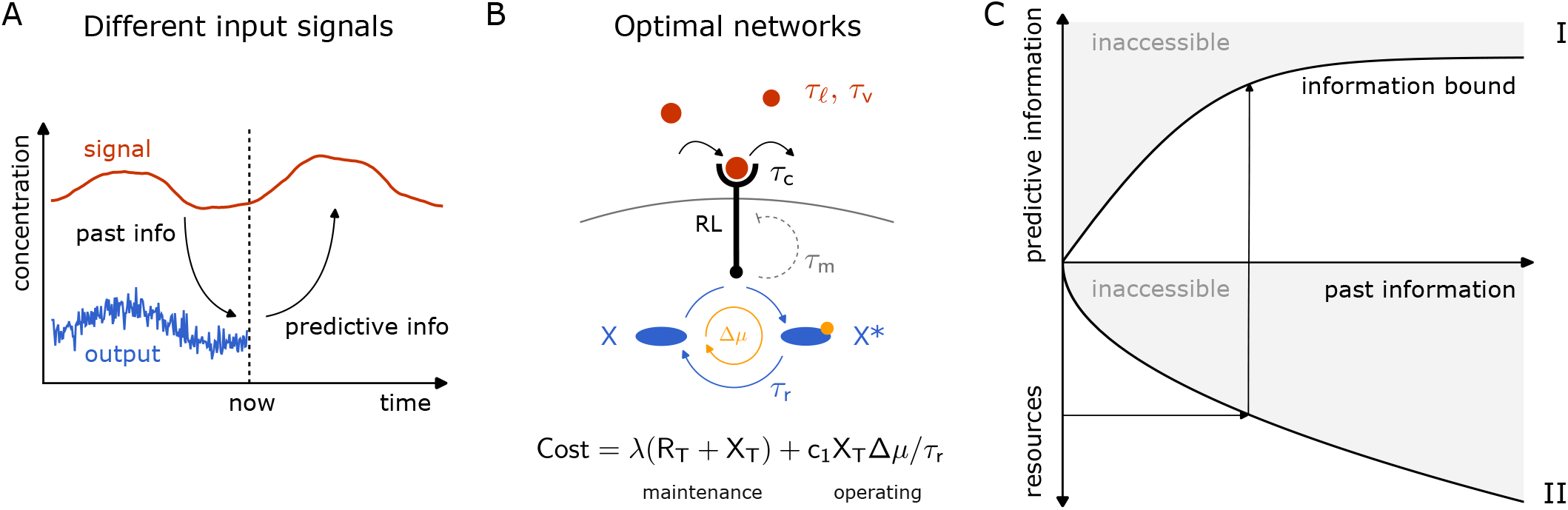
Cells use biochemical networks to remember the past and predict the future. (A) Cells compress the past input into the dynamics of the signalling network from which the future input is then predicted. (B) The optimal topology of the network for predicting the future signal depends on the temporal statistics of the input signal. Push-pull networks, consisting of chemical modification cycles or GTPase cycles, can optimally predict the future value of Markovian signals, with correlation time *τ*_*ℓ*_; derivative-taking networks, like the *E. coli* chemotaxis system, can optimally predict the future derivative of non-Markovian signals, with correlation time *τ_v_*. The push-pull network consists of a receptor that drives a downstream phosphorylation cycle. The ligand binds the receptor with a correlation time *τ*_c_. The push-pull network, driven by ATP turnover, integrates the receptor with an integration time *τ*_r_. The chemotaxis system is a push-pull network, yet augmented with negative feedback on the receptor activity via methylation on a timescale *τ*_m_, as indicated by the dashed grey line. The total resource cost consists of a maintenance cost of receptor and readout synthesis at the growth rate λ, and an operating cost of driving the cycle. (C) The predictive information on the future signal *I*_pred_ is fundamentally bounded by how much information *I*_past_ it has about the past signal (panel I), which in turn is limited by the resources necessary to build and operate the biochemical network (panel II) [6].

How close cells can come to this bound depends on the design of the intracellular biochemical network that senses and processes the environmental signals (Fig. 1B). To maximize the predictive power the cell must use its memory effectively: it should extract only those characteristics from the present and past signal that are most informative about the future [7]. Whether it can do so, is determined by the topology of the signaling network. Moreover, like any information processing device, biochemical networks require resources to be built and run. Molecular components are needed to construct the network, space is required to accommodate the components, time is needed to process the information, and energy is required to synthesize the components and operate the network [10]. These resources constrain the design and performance of any biochemical network, and the capacity to sense and process information is no exception (Fig. 1C-II).

Cellular signaling systems provide a unique opportunity for revealing the resource requirements for prediction. Cells live in a highly dynamic environment, with temporal statistics that are expected to vary markedly. Moreover, signaling networks have distinct topologies, which are likely tailored to the temporal statistics of the environment [7]. In addition, for cellular systems we can actually quantify the information processing capacity as a function of the resources that are necessary to build and run them—protein copies, time, and energy [10, 11].

Cellular systems are thus ideal for elucidating the relationships between future and past information, system design (i.e. network topology) and resource constraints. Here, we derive the bound on the prediction precision as set by the information extracted from the past signal for two types of input signals. We will determine how close cellular networks can come to this bound, and how this depends on the topology of the network and the resources to build and run it.

We find that for the two classes of input signals studied, cellular networks exists that can reach the information bound, yet reaching the bound is exceedingly costly. The first class of input signals consists of Markovian signals. Using the Information Bottleneck Method (IBM) [8, 12], we first show that the system that reaches the information bound copies the most recent input signal into the output from which the future input is predicted. Push-pull networks consisting of chemical modification or GTPase cycles, which are ubiquitous in prokaryotic and eukaryotic cells [13, 14], should be able to reach the information bound, because they are at heart copying devices [10, 11]. Yet, copying the most recent input into the output is extremely costly, because the operating cost, as set by the chemical power to drive the cycle, diverges at high copying speed. More surprisingly, our results show that the predictive and past information can be raised simultaneously by moving away from the information bound, even when the operating cost is negligible: the optimal system that maximizes the predictive information for a given protein synthesis cost is, in general, not at the information bound. The number of bits of past information per protein cost can be raised by increasing the integration time. While this decreases the predictive power per bit of past information, thereby moving the system away from the information bound, it can increase the total predictive information per protein cost. Our analysis thus highlights that not all bits of past information are equally costly, nor predictive.

Living cells that navigate their environment typically experience signals with persistence as generated by their own motion, which motivated us to study a simple class of non-Markovian signals. Moreover, these cells can typically detect changes in the concentration over a range of background concentrations that is orders of magnitude larger than the change in the concentration over the orientational correlation time of their movement. Our analysis reveals that in such a scenario the optimal kernel that allows the system to reach the information bound on predicting the future input derivative is a perfectively adaptive, derivative-taking kernel, precisely as the bacterium *E. coli* employs [15]. We again find, however, that reaching the information bound is prohibitively costly. The reason is that taking an instantaneous derivative, which is the characteristic of the input that is most informative about the future derivative, reduces the gain to zero because the system instantly adapts; the response becomes thwarted by biochemical noise. The optimal system that maximizes the predictive information under a resource constraint thus emerges from a trade-off between taking a derivative that is recent and one that is reliable. Finally, our analysis reveals that the *E. coli* chemotaxis system has been optimally designed to predict future concentration changes in shallow gradients.

## RESULTS

We focus on cellular signaling systems that respond linearly to changes in the input signal [11, 16–19]. These systems not only allow for analytical results, but also describe information transmission often remarkably well [19–22]. The output of these systems can be written as

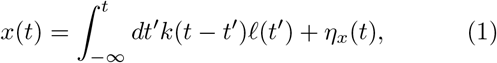

where *k*(*t*) is the linear response function, *ℓ*(*t*) the input signal, and *η_x_*(*t*) describes the noise in the output. We will consider stationary signals with different temporal correlations, obeying Gaussian statistics.

Any prediction about the future state of the environment must be based on information obtained from its past (Fig. 1C-I). In particular, the cell needs to predict the input *ℓ_τ_* ≡ *ℓ*(*t* + *τ*) at a time *τ* into the future from the current output *x_0_* ≡ *x*(*t*), which itself depends on the input signal in the past, ***L**_p_* ≡ (*ℓ*(*t*),*ℓ*(*t’*), …), with *t* > *t’* > …. The (qualitative) shape of the integration kernel *k*(*t*), e.g. exponential, adaptive or oscillatory, is determined by the topology of the signaling network [7]. The kernel shape describes how the past signal is mapped onto the current output, and hence which characteristics of the past signal the cell uses to predict the future signal. To maximize the accuracy of prediction, the cell should extract those features that are most informative about the future signal. These depend on the statistics of the input signal.

Deriving the upper bound on the predictive information as set by the past information is an optimisation problem, which can be solved using the IBM [8]. It entails the maximization of an objective function 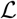:

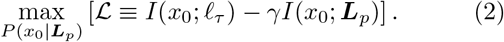

Here, *I_pred_* ≡ *I*(*x*_0_; *ℓ_τ_*) is the predictive information, which is the mutual information between the system’s current output xo and the future ligand concentration *ℓ_τ_*. The past information *I_past_* ≡ *I*(*x_0_*; ***L**_p_*) is the mutual information between *x*_0_ and the trajectory of past ligand concentrations ***L**_p_*. The Lagrange multiplier *γ* sets the relative cost of storing past over obtaining predictive information. Given a value of *γ*, the objective function in Eq. 2 is maximized by optimizing the conditional probability distribution of the output given the past input trajectory, *P*(*x*_0_|***L***_p_). For the linear systems considered here, this corresponds to optimizing the mapping of the past input signal onto the current output via the integration kernel *k*(*t*). Since our model obeys Gaussian statistics, we use the Gaussian IBM to derive the optimal kernel *k*^opt^(*t*) and the *information bound*, defined to be the maximum predictive information as set by the past information [12] (see Appendix C).

### Markovian signals

#### Optimal prediction of Markovian signals: biochemical copying

Arguably the most elementary type of signal, albeit perhaps the hardest to predict, is a Markovian signal. We consider a Markovian signal *ℓ*(*t*), of which the deviations 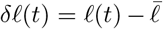 from its mean 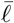 follow an Ornstein-Uhlenbeck (OU) process:

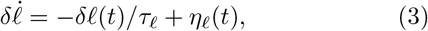

where *τ_ℓ_* is the correlation time of the fluctuations, and *ηℓ*(*t*) is Gaussian white noise, 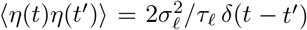, with 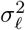 the amplitude of the signal fluctuations. This input signal obeys Gaussian statistics, characterized by 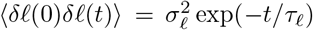. The optimal mapping is therefore a linear one. Utilizing the Gaussian IBM framework [12], we find that the optimal integration kernel is given by (see Appendix C2)

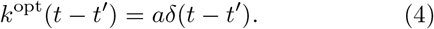

This optimal integration kernel corresponds to a signaling system that copies the current input into the output. This is intuitive, since for a Markovian signal there is no additional information in the past signal that is not already contained in the present one. The prefactor *a* determines the gain 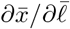 which together with the noise strength 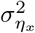 (Eq. 1) and the signal amplitude 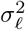 set the magnitude of the past and predictive information, *I*_past_ and *I*_pred_, respectively (see Appendix C1).

Fig. 2-I shows the maximum predictive information as set by the past information. This information bound applies to any linear system that needs to predict a Markovian signal. How close can biochemical systems come to this bound?

**FIG. 2.**
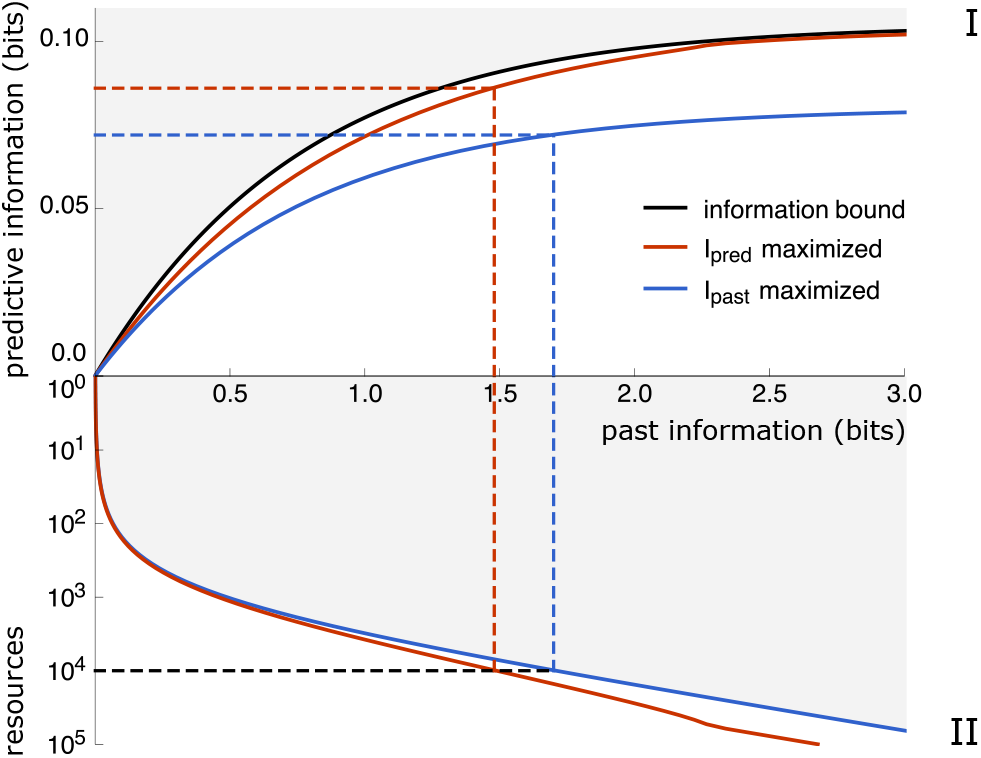
The optimal push-pull network is not at the information bound. Panel I: The black line is the information bound that maximizes the predictive information *I*_pred_ = *I*(*x*_0_;*ℓ_τ_*) for a given past information *I*_past_ = *I*(*x*_0_; ***L**_p_*). The red curve shows *I*_pred_ against *I*_past_ for systems in which *I*_pred_ has been maximized for a given resource cost *C* = *R_T_* + *X_T_*. The blue curve shows *I*_pred_ versus *I*_past_ for systems where *I*_past_ has been maximized for a given *C*. Panel II shows *I*_past_ against *C* for the corresponding systems. The forecast interval is *τ* = *τ_ℓ_*. The optimization parameters are the ratio *X*_T_/*R*_T_, *τ*_r_, *p* and *f* (see Appendix E). Parameter values: 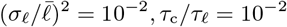,*τ*_c_/*τ*_*ℓ*_ = 10^-2^.

#### Push-pull network can be at the information bound, yet increase the predictive and past information by moving away from it

Although the upper bound on the accuracy of prediction is determined by the signal statistics, how close cells can come to this bound depends on the topology of the cellular signaling system, and the resources devoted to building and operating it. A network motif that could reach the information bound for Markovian signals is the push-pull network (Fig. 2), because it is at heart a copying device: it samples the input by copying the state of the input, e.g. the ligand-binding state of a receptor or the activation state of a kinase, into the activation state of the output, e.g. phosphorylation state of the readout [10, 11, 23].

We model the push-pull network in the linear-noise approximation:

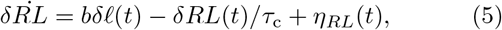

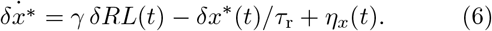

Here, *δRL* represents the number of ligand-bound receptors and *δx\** the number of modified readout molecules, defined as deviations from their mean values; *b* and *γ* are parameters that depend on the number of receptor and readout molecules, *R*_T_ and *X*_T_ respectively, the fraction of ligand-bound receptors *p* and active readout molecules *f*; *η_RL_* and *η_x_* are Gaussian white noise terms (see Appendix E). Key parameters are the correlation time of receptor-ligand binding, *τ*_c_, and the relaxation time of x*, *τ*_r_. The latter determines for how long *x*\* carries information on the ligand-binding state of the receptor and thus sets the integration time. The readout-modification dynamics yield an exponential integration kernel *k*(*t*) ∝ exp(*-t*/*τ*_r_), which in the limit *τ*_r_ → 0 reduces to a *δ*-function, hinting that the system may reach the information bound.

How much information cells can extract from the past signal depends on the resources devoted to building and operating the network (Fig. 2-II). We define the total resource cost to be:

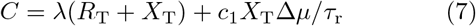

The first term expresses the fact that over the course of the cell cycle all components need to be duplicated, which means that they have to be synthesized at a speed that is at least the growth rate λ. The second term describes the chemical power that is necessary to run the push-pull network [10, 11]; it depends on the flux through the network, *X*_T_/*τ*_r_, and the free-energy drop Δ*μ* over a cycle, e.g. the free energy of ATP hydrolysis in the case of a phosphorylation cycle. The coefficient *c*_1_ describes the relative energetic cost of synthesising the components during the cell cycle versus that of running the system. For simplicity, we first consider the scenario that the cost is dominated by that of protein synthesis, setting *c*_1_ → 0. While in this scenario *R*_T_ + *X*_T_ is constrained, *X*_T_/*R*_T_ and other system parameters are free for optimization.

The available resources put a hard bound on the information *I*_past_ that can be extracted from the past signal, which in turn sets a hard limit on the predictive information *I*_pred_ (Fig. 1C). To maximize the predictive information, it therefore seems natural to maximize the past information *I*_past_ for a given resource cost *C*. The blue line in Fig. 2-II shows the result for the push-pull network. We then compute the corresponding predictive information for the systems along this line, which is the blue line in Fig. 2-I. Strikingly, the resulting information curve lies far below the information bound, i.e. the upper bound on the predictive information as set by the past information (black line, Fig. 2-I). This shows that systems that maximize past information under a resource constraint, do not in general also maximize predictive information. It implies that not all bits of past information are equally predictive about the future.

Precisely because not all bits of past information are equally predictive about the future, it is paramount to directly maximize the predictive information for a given resource cost in order to obtain the most efficient prediction device. This yields the red lines in panels I and II in Fig. 2. It can be seen that the predictive information is higher while the past information is lower, as compared to the information curves of the systems optimized for maximizing the past information under a resource constraint (blue lines). It reflects the idea that not all bits are equally predictive. More surprisingly, while the bound on the predictive information as set by the resource cost (red line panel I) is close to the bound on the predictive information as set by the past information (black line), it does remain lower. This is surprising, because the push-pull network is a copying device [10, 23], which can, as we will also show below, reach the latter bound. These two observations together imply that not all bits of past information are equally costly. If they were, the cell would select under the two constraints the same bits based on their predictive information content, and the bound on the predictive information as set by the resource cost would overlap with that as set by the past information.

We thus find that not all bits of past information are equally predictive, nor equally costly. As we show next, it implies that the optimal information processing system faces a trade-off between using those bits of past information that are most informative about the future and those that are cheapest.

#### Trade-off between cost and predictive power per bit

To understand the connection between predictive and past information, and resource cost, we map out the region in the information plane that can be reached given a resource constraint *C* (Fig. 3A, green region). We immediately make two observations. Firstly, the system can indeed reach the information bound. Secondly, the system can increase both the past and the predictive information by moving away from the bound. To elucidate these two observations, we investigate the system along the isocost line of *C* = 10^4^, which together with the information bound envelopes the accessible region for the maximum resource cost *C* ≤ 10^4^.

**FIG. 3.**
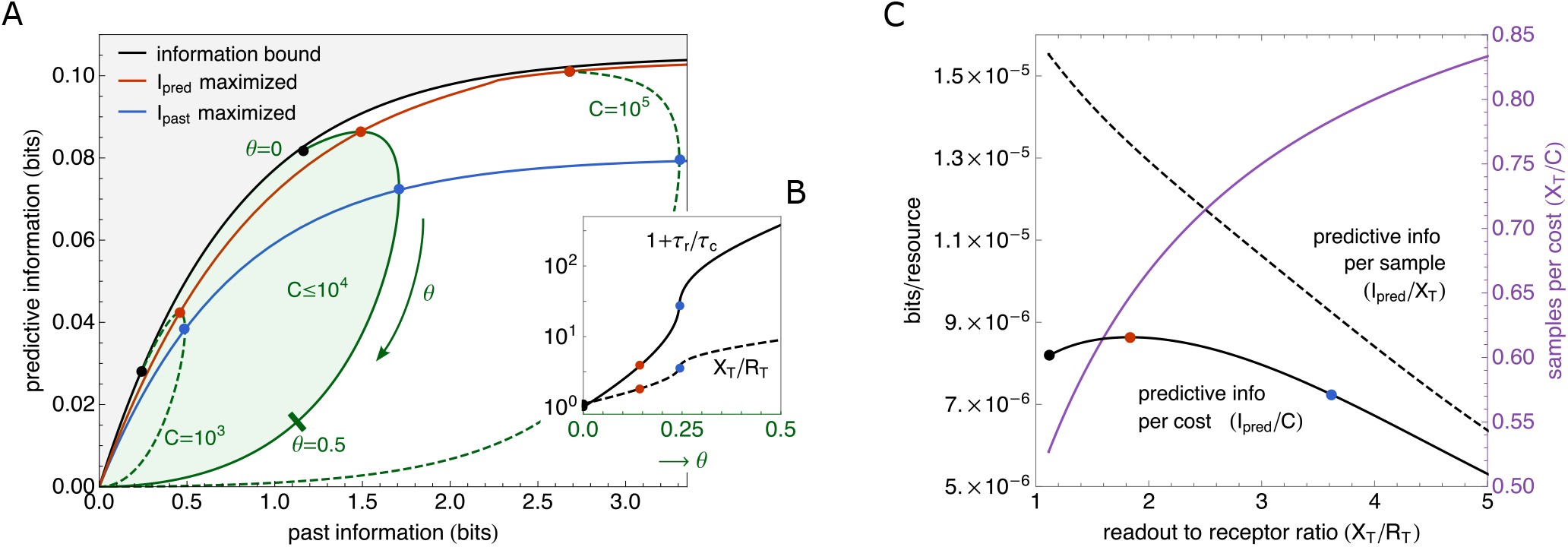
The push-pull network maximizes the predictive power under a resource constraint by moving away from the information bound. (A) The region of accessible predictive information *I*_pred_ = *I*(*x*_0_;*ℓ*_*τ*_) and past information *I*_past_ = *I*(*x*_0_; ***L**_p_*) in the push-pull network under a resource constraint *C* ≤ (*R*_T_ + *X*_T_), for the Markovian signals specified by Eq. 3 (green). The black line is the information bound at which *I*_pred_ is maximized for a given *I*_past_. The push-pull network can be at the information bound (black points), but maximizing *I*_pred_ for a resource constraint *C* moves the system away from it. The red and blue lines connect, respectively, the points where *I*_pred_ and *I*_past_ are maximized along the green isocost lines (the contourlines of constant *C*); they correspond to the red and blue lines in Fig. 2, respectively. The accessible region of *I*_pred_ and *I*_past_ for a given *C* has been obtained by optimizing over *τ*_r_, *p*, *f*, and *X*_T_/*R*_T_. The forecast interval is *τ* = *τ*_*ℓ*_. (B) The integration time *τ*_r_ over the receptor correlation time *τ*_c_, *τ*_r_/*τ*_c_, and the ratio of the number of readout and receptor molecules, *X*_T_/*R*_T_, as a function of the distance *θ* along the isocost line corresponding to *C* = 10^4^ in panel A; the red and blue points denote where *I*_pred_ and *I*_past_ are maximized along the contourline, respectively. For *θ* → 0, *τ*_r_ → 0: the system is an instantaneous responder, which is essentially at the information boundary; as predicted by the optimal resource allocation principle, *X*_*τ*_ = *R*_*τ*_. The system can increase *I*_pred_ and *I*_past_ by increasing *τ*_r_ and *X*_T_/*R*_T_. (C) While this decreases the predictive information *I*_pred_ per physical bit of past information, *I*_pred_/*X*_T_ (dashed line), increasing *X*_T_/*R*_T_ does increase the number of physical bits per resource cost, *X*_T_/*C* (purple line). This trade-off gives rise to an optimal predictive information per resource cost, *I*_pred_/*C* (red dot on solid black line). Parameter values unless specified: 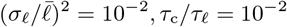,*τ*_c_/*τ*_ℓ_ = 10^-2^.

Along the isocost line, the ratio of the number of readout over receptor molecules is 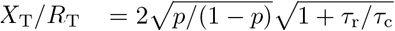 (see Appendix E3). This can be understood intuitively using the optimal resource allocation principle [10]. It states that in a sensing system that employs its proteins optimally, the total number of independent concentration measurements at the level of the receptor during the integration time *τ*_r_, *R*_T_(1 + *τ*_r_/*τ*_c_), equals the number of readout molecules *X*_T_ that store these measurements, so that neither the receptors nor the readout molecules are in excess. This design principle specifies, for a given integration time *τ*_r_, the ratio *X*_T_/*R*_T_ at which the readout molecules sample each receptor molecule roughly once every receptor correlation time *τ*_c_.

While the optimal allocation principle gives the optimal ratio *X*_T_/*R*_T_ of the number of readouts over receptors for a *given* integration time *τ*_r_, it does not prescribe what the optimal integration time *τ*_r_^opt^, and hence (globally) optimal ratio 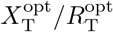, is that maximizes *I*_pred_ for a given resource constraint *C* = *R*_T_ + *X*_T_. Fig. 3B shows that as the distance *θ* along the isocost line is increased, *τ*_r_ and hence *X*_T_/*R*_T_ increase monotonically. Near the information bound, corresponding to *θ* = 0, the integration time *τ*^r^ is zero and the number of readout molecules equals the number of receptor molecules: *X*T = *R*T. In this limit, the push-pull network is an instantaneous responder, with an integration kernel given by Eq. 4; only the finite receptor correlation time *τ*_c_ prevents the system from fully reaching the information bound. Yet, as *θ* increases and the system moves away from the bound, the predictive and past information first rise along the contour, and thus with *X*T/*R*T and *τ*_r_, before they eventually both fall.

To understand why the predictive and past information first rise and then fall with *X*_T_/*R*_T_ and *τ*_r_, we note that each readout molecule constitutes 1 physical bit and that its binary state (phosphorylated or not) encodes at most 1 bit of information on the ligand concentration. The number of readout molecules *X*_T_ thus sets a hard upper bound on the sensing precision and hence the predictive information. To raise this bound, *X*_T_ must be increased. For a given resource constraint *C* = *R*_T_ + *X*_T_, *X*_T_ can only be increased if the number of receptors *R*_T_ is simultaneously decreased. However, the cell infers the concentration not from the readout molecules directly, but via the receptor molecules: a readout molecule is a sample of the receptor that provides at most 1 bit of information about the ligand-binding state of a receptor molecule, which in turn provides at most 1 bit of information about the input signal. To raise the lower bound on the predictive information, the information on the input must increase at both the receptor and the readout level.

To elucidate how this can be achieved, we note that the maximum number of independent receptor samples and hence concentration measurements is given by 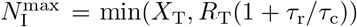 [10]. For *θ* > 0, the system can increase 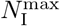 if, and only if, *X*_T_ and *R*_T_(1 + *τ*_r_/*τ*_c_) can be raised simultaneously. This can be achieved, while obeying the constraint *C* = *X*_T_ + *R*_T_, by decreasing *R*_T_ yet increasing *τ*_r_ (Fig. 3B). This is the mechanism of time averaging, which makes it possible to increase the number of independent receptor samples [11], and explains why both the predictive and the past information initially increase (Fig. 3C). However, as *τ*_r_ is raised further, the receptor samples become older: the readout molecules increasingly reflect receptor states in the past that are less informative about the future ligand concentration. The collected bits of past information have become less predictive about the future (Fig. 3C). For a given resource cost, the cell thus faces a trade-off between maximizing the number of physical bits of past information (i.e. the receptor samples *X*_T_) and the predictive information per bit. This antagonism gives rise to an optimal integration time 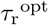 that maximizes the total predictive information *I*_pred_ (Fig. 3C).

Interestingly, while *I*_pred_ decreases beyond 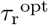, the past information *I*_past_ first continues to rise because 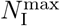 still increases. However, when the integration time becomes longer than the input signal correlation time, the correlation between input and output will be lost and *I*_past_ will fall too.

#### Chemical power prevents the system from reaching the information bound

So far, we have only considered the cost of maintaining the cellular system, the protein cost *C* = *R*_T_ + *X*_T_. Yet, running a push-pull network also requires energy. As Eq. 7 shows, the running cost scales with the flux around the phosphorylation cycle, which is proportional to the inverse of the integration time, **τ*_r_^-1^*. The power thus diverges for *τ*_r_ → 0. Since the information bound is reached precisely in this limit, it is clear that the chemical power prevents the push-pull network from reaching the bound (see Fig. 3 in the appendix).

### Non-Markovian signals

#### Predicting the future change

The push-pull network can optimally predict Markovian signals, yet not all signals are expected to be Markovian. Especially organisms that navigate through an environment with directional persistence will sense a non-Markovian signal, as generated by their own motion. Moreover, when these organisms need to climb a concentration gradient, as *E. coli* during chemotaxis, then knowing the change in the concentration is arguably more useful than knowing the concentration itself. Indeed, it is well known that the kernel of the *E. coli* chemotaxis system detects the (relative) change in the ligand concentration by taking a temporal derivative of the concentration [15]. However, as we will show here, the converse statement is more subtle. If the system needs to predict the (future) change in the signal, then the optimal kernel is not necessarily one that is based on the derivative only: in general, the optimal kernel uses a combination of the signal value and its derivative. However, the *E. coli* chemotaxis system can respond to concentrations that vary between the dissociation constants of the inactive and active state of the receptors, which differ by several orders of magnitude [24]. This range of possible background concentrations is much larger than the typical concentration change over the orientational correlation time of the bacterium. As our analysis below reveals, in this regime the optimal kernel is a perfectly adaptive, derivative-taking kernel that is insensitive to the current signal value, precisely like that of the *E. coli* chemotaxis system [15, 25–28]. Our analysis thus predicts that this system has an adaptive kernel, because this is the optimal kernel for predicting concentration derivatives over a broad range of background concentrations.

To reveal the signal characteristics that control the shape of the optimal integration kernel, we will consider the family of signals that are generated by a harmonic oscillator:

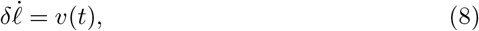

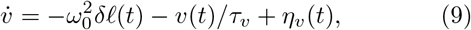

where *δℓ* is the deviation of ligand concentration from its mean 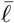, *v* its derivative, *τ_v_* a relaxation time, *η_v_* a Gaussian white noise term, and the frequency 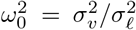 controls the variance 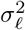 of the concentration and that of its derivative 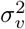.

Using the IBM framework it can be shown that the optimal encoding that allows the system to reach the information bound, is based on a linear combination of the current concentration *ℓ*(*t*) and its derivative *v*(*t*), such that the output *x*(*t*) is given by (Appendix C3):

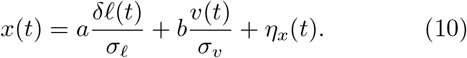

This can be understood by noting that while the signal of Eqs. 8 and 9 is non-Markovian in the space of *ℓ*, it is Markovian in *ℓ* and *v*: all the information on the future signal is thus contained in the current concentration and its derivative. To maximize the predictive information *I*_pred_ = *I*(*x*_0_; *v*_*τ*_) between the current output *x*_0_ and the future derivative of the input *v*_*τ*_ for a given amount of past information *I*_past_ = *I*(*x*_0_; ***L**_p_*), i.e to reach the information bound for predicting the future signal derivative, the coefficients must obey

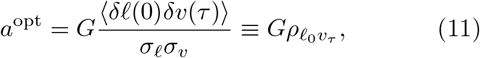

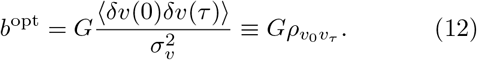

Here, *G* is the gain, which together with the noise 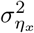 sets the scale of I_pred_ and *I*_past_, *ρ*_*ℓ*_0_*v*_*τ*__ is the cross-correlation coefficient between the current concentration value *ℓ*_0_ and the future concentration derivative *v*_*τ*_ and *ρ*_*v*_0_*v*_*τ*__ that between the current and future derivative (Appendix C3). These expressions can be understood intuitively: if the future signal derivative that needs to be predicted is correlated with the current signal derivative, it is useful to include in the prediction strategy the current signal derivative, leading to a non-zero value of *b*^opt^. Perhaps more surprisingly, if the future signal derivative is also correlated with the current signal *value*, then the system can enhance the prediction accuracy by also including the current signal value, yielding a non-zero *a*^opt^. Clearly, in general, to optimally predict the future signal change, the system should base its prediction on both the current signal value and its derivative.

The degree to which the systems bases its prediction on the current value versus the current derivative depends on the relative magnitudes of *a*^opt^ and *b*^opt^, respectively. In Appendix B2, we show that when the concentration change over the timescale *τ*_v_, *σ*_v_ *τ*_v_, is much smaller than the range of possible concentrations *σ_ℓ_* that the bacterium can experience, i.e. when σ_v_*τ*_v_ ≪ *σ_ℓ_* such that 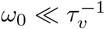, the cross-correlation coefficient *ρ*_*ℓ*_o_*v*_*τ*__ vanishes, such that *a*^opt^ becomes zero (see Eq. 11). The optimal kernel has become a perfectly adaptive, derivative-taking kernel. We emphasize that while we have derived this result for the class of signals defined by Eqs. 8 and 9, the idea is far more generic. In particular, while we do not know the temporal structure of the ligand statistics that *E. coli* experiences, we do know that it can detect concentration changes over a range of background concentrations that is much wider that the typical concentration change over a run, such that the correlation between the concentration value and its future change is likely to be very small. As our analysis shows, a perfectively adaptive kernel then emerges naturally from the requirement to predict the future concentration change.

While the class of signals specified by Eqs. 8 and 9 is arguably limited, it does describe the biologically important regime of chemotaxis in shallow gradients. In the limit that *ω*_0_ ≪ *τ*_v_^-1^, Eq. 9 reduces to 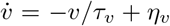. In shallow gradients, the stimulus only weakly affects the swimming behavior, such that the perceived signal is mostly determined by the intrinsic orientational dynamics of the bacterium in the absence of a gradient. In this regime, the temporal statistics of the concentration derivative *v* is completely determined by the steepness of the concentration gradient *g* and the swimming statistics of the bacterium in the absence of a gradient:

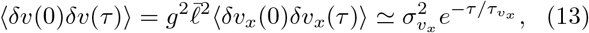

where the latter is the autocorrelation function of the (positional) velocity of the bacterium in the absence of a gradient. It is a characteristic of the bacterium, not of the environment, and has been measured to decay exponentially with a correlation time *τ*_*v*__*x*_ [18], precisely as our model, with *τ*_v_ = *τ*_*v*__*x*_, predicts. This correlation time is on the order of the typical run time of the bacterium in the absence of a gradient, *τ*_*v*_ ~ 0.9s [18].

#### Finite resources prevent the chemotaxis system from taking an instantaneous derivative and reaching the information bound

The above analysis indicates that the chemotaxis system seems ideally designed to predict the future concentration change, because its integration kernel is nearly perfectly adaptive [15, 25–28]. But how close can this system come to the information bound for the non-Markovian signals specified by Eqs. 8 and 9?

To address this, we consider a molecular model that can accurately describe the response of the chemotaxis system to a wide range of time-varying signals [29–32]. In this model, the receptors are partitioned into clusters. Each cluster is described via a Monod-Wyman-Changeux model [33]. While each receptor can switch between an active and an inactive conformational state, the energetic cost of having different conformations in the same cluster is prohibitively large. Each cluster is thus either active or inactive. Ligand binding favors the inactive state while methylation does the opposite. Lastly, active receptor clusters can via the associated kinase CheA phosphorylate the downstream messenger protein CheY.

Linearizing around the steady state, we obtain:

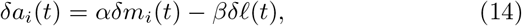

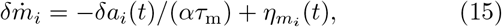

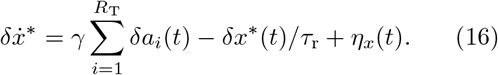

Here, *δ_a__i_* (*t*) and *δ_m__i_*(*t*) are the deviations of the activity and methylation level of receptor cluster *i* from their steady-state values, and *R*_T_ is the total number of receptor clusters; *δℓ*(*t*) and *δ_x_*\*(*t*) are, respectively, the deviations of the ligand and CheYp concentration from their steady-state values; *τ*_m_ and *τ*_r_ are the timescales of receptor methylation and CheY_p_ dephosphorylation; *η*_m__*i*_ and *η_x_* are independent Gaussian white noise sources. In Eq. 14, we have assumed that ligand binding is much faster than the other timescales in the system, so that it can be integrated out. There is therefore no need to time average receptor-ligand binding noise, which means that, in the absence of running costs, the optimal receptor integration time *τ*_r_ is zero. In what follows, we set *τ*_r_ to the value measured experimentally, *τ*_r_ ≈ 100ms [10, 34]. We consider the non-Markovian signals specified by Eqs. 8 and 9 in the physiologically relevant limit *ω*_0_ *→* 0, such that the optimal kernel is perfectly adaptive, like that of *E. coli*. For these signals, we determine the accessible region of *I*_past_ and *I*_pred_ under a resource constraint *C* = *R*_T_ + *X*_T_ (see Fig. 4) by optimizing over the methylation time *τ*_m_ and the ratio of readout over receptor molecules *X*_T_/*R*_T_. The forecast interval *τ* is set to *τ_v_*, but we emphasize that the optimal design is independent of the value of *τ* (see Appendix F4).

**FIG. 4.**
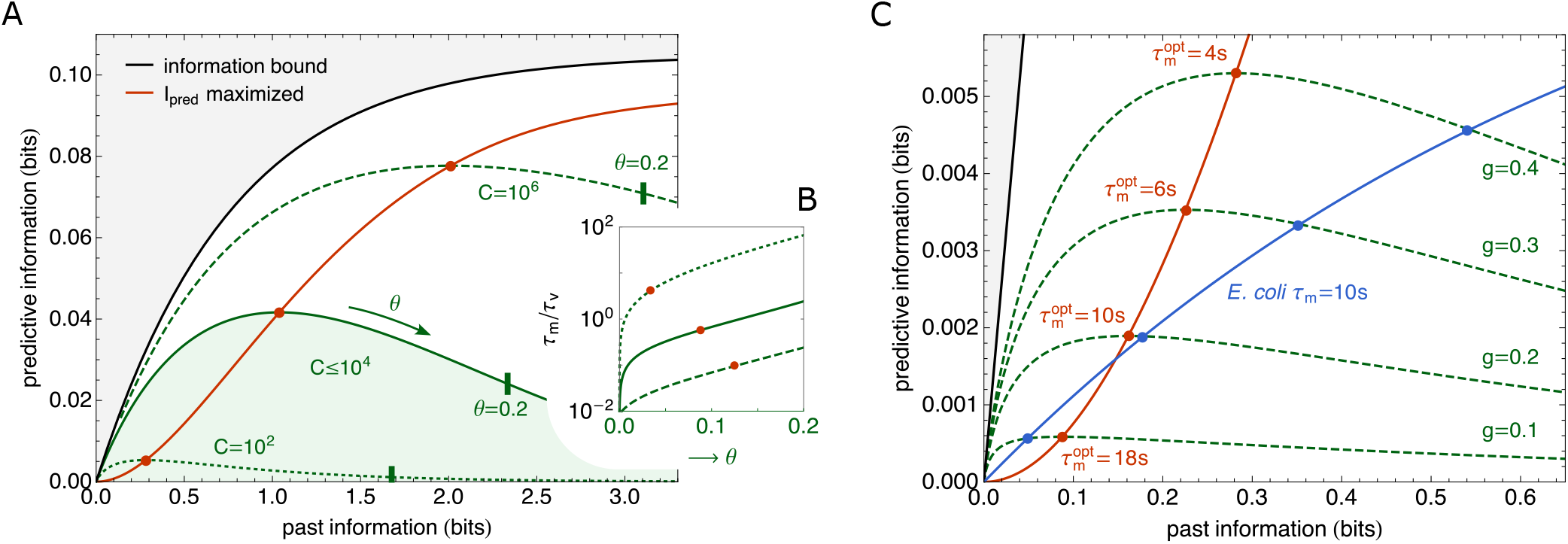
Finite resources prevent chemotaxis system from reaching the information bound. (A) The region of accessible predictive information *I*_pred_ = *I*(*x*_0_;*v*_*τ*_) and past information *I*_past_ = *I*(*x*; ***L**_p_*) for the chemotaxis system under a resource constraint *C* = **R*_T_* + **X*_T_*, for the non-Markovian signals specified by Eqs. 8 and 9 (green). The black line shows the information bound at which *I*_pred_ is maximized for a given *I*_past_. The chemotaxis system is not at the information bound, but it does move towards it as *C* is increased. The red line connects the red points where *I*_pred_ is maximized for a given resource cost *C*. The accessible region of *I*_pred_ and *I*_past_ under a given resource constraint *C* = *R*_T_ + *X*_T_ is obtained by optimizing over the methylation time *τ*_m_ and the ratio of readout over receptor molecules *X*_T_/*R*_T_. The forecast interval is *τ* = *τ_v_*. (B) The methylation time *τ*_m_ over the input correlation time *τ_v_* as a function of the distance *θ* along the three respective isocost lines shown in panel A. The methylation time *τ*_m_ increases along the isocost line, but there exists an optimal *τ*_m_ that maximizes the predictive information, marked by the red points; *θ* → 0 corresponds to the origin of panel A, (*I*_pred_, *I*_past_) = (0, 0); the points where *θ* = 0.2 along the isocost lines of panel A are marked with a bar. As the resource constraint is relaxed (higher *C*), the optimal *τ*_m_ decreases: the system moves towards the information bound, where it takes an instantaneous derivative, corresponding to *τ*_r_,*τ*_m_ → 0. (C) The contourlines of *I*_pred_ and *I*_past_ for increasing values of the steepness *g* of an exponential ligand concentration gradient *ℓ*(*x*) = *ℓ*_0_^*e*^^*gx*^, keeping the total resource cost fixed at *C* = *R*_T_ + *X*_T_ = 10^4^; *τ*_m_ and *X*_T_/*R*_T_ have been optimized. It is seen that the maximal predictive information *I*_pred_ under the resource constraint *C* (marked by the red points) increases with the gradient steepness. The blue line shows *I*_pred_ and *I*_past_ for the *E. coli* chemotaxis system with *τ*_m_ = 10s and *X*_T_ = *R*_T_ = 5000 fixed at their measured values [35]. Our analysis predicts that this system has been optimized to detect shallow gradients. Parameter values unless specified: *τ*_r_ = 100ms [10, 34]; *τ*_v_ = 0.9s and 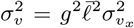, with 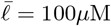 and 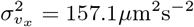 [18]; *ω*_0_ → 0; *g* is given in units of mm^-1^; in A, g = 4/mm.

Fig. 4A shows that the chemotaxis system is, in general, not at the information bound that maximizes the predictive information *I*_pred_ = *I*(*x*_0_; *v_τ_*) for a given past information *I*_past_ = *I*(*x*_0_; ***L**_p_*). The optimal systems that maximize *I*_pred_ under a resource constraint *C*, marked by the red dots, are indeed markedly away from the information bound. Yet, as the resource constraint is relaxed and *C* is increased, the optimal system moves towards the bound. Panel B shows that the methylation time *τ*_m_ rises along the three respective isocost lines of panel A. It highlights that there exists an optimal methylation time 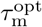 that maximizes the predictive information *I*_pred_. Moreover, 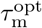 decreases as the resource constraint is relaxed. Along the respective isocost lines, *X*_T_/*R*_T_ varies only mildly (see Fig. 5 in the appendix).

These observations can be understood by noting that the system faces a trade-off between taking a derivative that is recent versus one that is robust. All the information on the future derivative, which the cell aims to predict, is contained in the current derivative of the signal; measuring the current derivative would allow the system to reach the information bound. However, computing the recent derivative is extremely costly. The cell takes the temporal derivative of the ligand concentration at the level of the receptor via two antagonistic reactions that occur on two distinct timescales: ligand binding rapidly deactivates the receptor, while methylation slowly reactivates it [30]. The receptor ligand-occupancy thus encodes the current concentration, the methylation level stores the average concentration over the past *τ*_m_, and the receptor activity reflects the difference between the two—the temporal derivative of the signal over the timescale *τ*_m_. To obtain an instantaneous derivative, *τ*_m_ must go to zero. However, this dramatically reduces the gain; in fact, in this limit, the gain is zero, because the receptor activity instantly adapts to the change in the ligand concentration. Since the push-pull network downstream of the receptor is a device that samples the receptor stochastically [10, 36], the gain, i.e. the change in the receptor activity due to the signal, must be raised to lift the signal above the sampling noise. This requires a finite methylation time *τ*_m_: as we show in Appendix F3, the gain increases monotonically with *τ*_m_. The trade-off between a recent derivative and a reliable one gives rise to an optimal methylation time 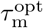 that maximizes the predictive information for a given resource cost.

The same analysis also explains why the optimal methylation time 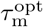 decreases and the predictive information increases when the resource constraint is relaxed. The sampling noise in estimating the average receptor activity decreases as the number of readout molecules increases [10, 36]. A smaller gain is thus required to lift the signal above the sampling noise. In addition, a larger number of receptors decreases the noise in the methylation level, which also allows for a smaller gain, and hence a smaller methylation time. These two effects together explain why 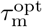 decreases and *I*_pred_ increases with *C*.

Fig. 4A also shows that the past information *I*_past_ = *I*(*x*_0_; ***L**_p_*) does not return to zero along the contourline of constant resource cost. Along the contourline, the methylation time *τ*_m_ rises (Fig. 4B). While the predictive information *I*_pred_ exhibits an optimal methylation time *τ*_m_^opt^, the past information *I*_past_ continues to rise with *τ*_m_ because the system increasingly becomes a copying device, rather than one that takes a temporal derivative.

#### Comparison with experiment

To test our theory, we study the predictive power of the *E. coli* chemotaxis system as a function of the steepness of the ligand concentration gradient, keeping the resource constraint at the biologically relevant value of *C* = *R*_T_ + *X*_T_ = 10^4^ [35]. Panel *C* of Fig. 4 shows *I*_pred_ and *I*_past_ for cells swimming in an exponential concentration gradient *ℓ*(*x*) = *ℓ*_0*e*_^*gx*^, for different values of the gradient steepness *g*; along the green iso-steepness lines *τ*_m_ is varied and *X*_T_/*R*_T_ is optimized to maximize *I*_pred_ and *I*_past_, with the red dots marking 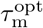, while along the blue line *τ*_m_ and *X*_T_ and *R*_T_ are fixed at their experimentally measured values [29, 30, 35]. Clearly, both the predictive and the past information rise as the gradient steepness *g* increases—a steeper concentration gradient yields a larger change in the concentration, and thus a stronger signal.

More interestingly, in the optimal system I_pred_ rises much faster with *I*_past_ (red line) than in the *E. coli* system (blue line). A steeper gradient g yields a stronger input signal, which raises the signal above the sampling noise more. This allows the optimal system to take a more recent derivative, with a smaller *τ*_m_, which is more informative about the future. In contrast, the methylation time *τ*_m_ of the *E. coli* chemotaxis system is fixed. As Fig. 4C shows, this value is beneficial for detecting shallow gradients, 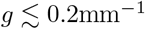. Moreover, in this regime, not only I_pred_ but also *I*_past_ are close to the respective values for the optimal system. For steeper gradients *I*_past_ becomes much higher in the *E. coli* system than in the optimal one, even though *I*_pred_ remains lower. The bacterium increasingly collects information that is less informative about the future. Taken together, these results strongly suggest that the system has been optimized to predict future concentration changes in shallow gradients, which necessitate a relatively long methylation time.

## DISCUSSION

Cellular systems need to predict the future signal by capitalizing on information that is contained in the past signal. To this end, they need to encode the past signal into the dynamics of the intracellular biochemical network from which the future input is inferred. To maximize the predictive information for a given amount of information that is extracted, the cell should store those signal characteristics that are most informative about the future signal. For a Markovian signal obeying an Ornstein-Uhlenbeck process this is the current signal value, while for the non-Markovian signal corresponding to an underdamped particle in a harmonic well, this is the current signal value and its derivative. As we have seen here, cellular systems are able to extract these signal characteristics: the push-pull network can copy the current input into the output, while the chemotaxis network can take an instantaneous derivative. We have thus demonstrated that at least for two classes of signals, cellular systems are in principle able to extract the most predictive information, allowing them to reach the information bound.

Yet, our analysis also shows that extracting the most relevant information can be exceedingly costly. To copy the most recent input signal into the output, the integration time of the push-pull network needs to go to zero, which means that the chemical power diverges. Moreover, taking an instantaneous derivative reduces the gain to zero, such that the signal is no longer lifted above the inevitable intrinsic biochemical noise of the signalling system. In fact, taking the chemical power cost to drive the adaptation cycle into account [27, 37] would push the system away from the information bound even more.

While information is a resource—the cell cannot predict the future without extracting information from the past signal—the principal resources that have a direct cost are time, building blocks and energy. The predictive information per protein and energy cost is therefore most likely a more relevant fitness measure than the predictive information per past information. Our analysis reveals that, in general, it is not optimal to operate at the information bound: cells can increase the predictive information for a given resource constraint by moving away from the bound. Increasing the integration time in the push-pull network reduces the chemical power and makes it possible to take more concentration measurements per protein copy. And increasing the methylation time in the chemotaxis system increases the gain. Both enable the system to extract more information from the past signal. Yet, increasing the integration time or the methylation time also means that the information that has been collected, is less informative about the future signal. This interplay gives rise to an optimal integration and methylation time, which maximize the predictive information for a given resource constraint. This argument also explains why the respective systems move towards the information bound when the resource constraint is relaxed: Increasing the number of receptor and readout molecules allows the system to take more instantaneous concentration measurements, which makes time averaging less important, thus reducing the integration time.

Increasing the number of readout molecules also reduces the error in sampling the receptor state. This makes it easier to detect a change in the receptor activity resulting from the signal, thus allowing for a smaller dynamical gain and a shorter methylation time.

Information theory shows that the amount of transmitted information depends not only on the characteristics of the information processing system, but also on the statistics of the input signal. While much progress has been made in characterizing cellular signalling systems, the statistics of the input signal is typically not known, with a few notable exceptions [38]. Here, we have focussed on two classes of input signals, but it seems likely that the signals encountered by natural systems are much more diverse. It will be interesting to extend our analysis to signals with a richer temporal structure [9], and see whether cellular systems exist that can optimally encode these signals for prediction.

Finally, while we have analyzed the design of cellular signaling networks to optimally predict future signals, we have not addressed the utility of information for function or behavior. It is clear that many functional or behavioral tasks, like chemotaxis [18], require information, but what the relevant bits of information are is poorly understood [7]. Moreover, cells ultimately employ their resources— protein copies, time, and energy—for function or behavior, not for processing information per se. Here, we have shown that maximizing predictive information under a resource constraint, *C* → *I*_past_ → *I*_pred_, does not necessarily imply maximizing past information. This hints that optimizing a functional or behavioral task under a resource constraint, *C* → *I*_pred_ → function, may not imply maximizing the predictive information necessary to carry out this task.

## ACKNOWLEDGMENTS

We thank Jenny Poulton, Manuel Reinhardt, Michael Vennettilli and Daan de Groot for many useful discussions. This work is part of the Dutch Research Council (NWO) and was performed at the research institute AMOLF. This project has received funding from the European Research Council (ERC) under the European Union’s Horizon 2020 research and innovation program (grant agreement No. 885065).

