## Appendices for "Trade-offs between cost and information in cellular prediction"

(Dated: January 10, 2023)

### CONTENTS

A. General 1
  1. Linear signalling networks 1
  2. Integration kernels, power spectra, and correlation functions 2

B. Signals and statistics 3
  1. Markovian signal 3
  2. Non-Markovian signal 3

C. Information bottleneck framework and solutions 4
  1. Gaussian information bottleneck 4
  2. Markovian signal 6
  3. Non-Markovian signal 7

D. Past and predictive information for linear signalling networks 9

E. Push-pull network 10
  1. Model statistics 11
  2. Past and predictive information of the push-pull network 12
  3. Optimal resource allocation 13
  4. Operating costs diverge when approaching the information bound 13

F. Chemotaxis network 14
  1. Linear dynamics 16
  2. Parameter values 16
  3. Model statistics 16
  4. Past and predictive information of the chemotaxis network 19
  5. Optimal allocation 19

References 20

### Appendix A: General

#### 1. Linear signalling networks

Since the systems studied in the main text have a single steady state, we will study them in the linear-noise approximation [1]. For non-linear systems, the quality of the approximation improves with system size, but it can already be remarkably good for systems with only 10 copies [2–4]. In the linear-noise approximation, we expand the rate equations to first order around the steady state of the mean-field chemical rate equations, and compute the

*

noise at this steady state. In this approximation the network dynamics are a multidimensional Ornstein-Uhlenbeck (OU-)process:

$$\dot{\boldsymbol{\delta y}} = \mathcal{G}\boldsymbol{\delta s}(t) + \mathcal{J}\boldsymbol{\delta y}(t) + \mathcal{B}\boldsymbol{\xi}(t), \tag{A1}$$

where $\boldsymbol{\delta s}(t)$ is a length $k$ vector of input signals and $\boldsymbol{\delta y}$ is the vector of all network species of length $n$, both defined in terms of deviations from their mean. The vector $\boldsymbol{\xi}(t)$ describes the $m$ independent white noise processes associated with the $m$ network reactions; they have zero mean, unit variance, and are delta correlated: $\langle \xi_i(t)\rangle = 0$, $\langle \xi_i(t)\xi_j(t')\rangle = \delta_{ij}\delta(t-t')$, with $\delta_{ij}$ the Kronecker delta. The $n \times n$ matrix $\mathcal{J}$ is the Jacobian of the network, the $n \times k$ signal gain matrix $\mathcal{G}$ describes the strength by which each signal impacts each species directly, the $n \times m$ matrix $\mathcal{B}$ contains the noise strengths. The eigenvalues of the Jacobian $\mathcal{J}$ must be negative for the system to be stable, and we require all signals to be stationary.

#### 2. Integration kernels, power spectra, and correlation functions

We continue by deriving the stationary auto-correlation matrix of a multidimensional OU-process, such as Eq. A1, via the networks' power spectra. The power spectrum of a real-valued random process $X(t)$ is the squared modulus of its Fourier transform: $S_x(\omega) = \langle \delta\tilde{x}(-\omega)\delta\tilde{x}(\omega)\rangle$ and $S_{x\to y}(\omega) = \langle \delta\tilde{x}(-\omega)\delta\tilde{y}(\omega)\rangle$. Throughout this work we use the following conventions for the Fourier transform and its inverse: $\mathcal{F}\{f(t)\} \equiv \tilde{f}(\omega) = \int_{-\infty}^{\infty} dt f(t)\exp(-i\omega t)$ and $\mathcal{F}^{-1}\{\tilde{f}(\omega)\} = 1/(2\pi)\int_{-\infty}^{\infty} d\omega \tilde{f}(\omega)\exp(i\omega t) = f(t)$. To obtain the correlation functions from the power spectra we invoke the Wiener-Khinchin theorem.

The general solution to Eq. A1 is

$$\boldsymbol{\delta y}(t) = \int_{-\infty}^{t} dt'\, e^{\mathcal{J}(t-t')}\left(\mathcal{G}\boldsymbol{\delta s}(t') + \mathcal{B}\boldsymbol{\xi}(t')\right), \tag{A2}$$

which shows the two contributions to the time dependent solution: that of the external signal and that of the internal noise. The $n \times k$ matrix $e^{\mathcal{J}(t-t')}\mathcal{G}$ contains the integration kernels, its $(i,\, j)^{\text{th}}$ entry determines how the $j^{\text{th}}$ signal affects the $i^{\text{th}}$ system component over time. The $n \times m$ matrix $e^{\mathcal{J}(t-t')}\mathcal{B}$ is similar, but contains the functions that map the noise terms onto the system components. These matrices can be obtained by taking the Fourier transform of Eq. A1 and solving for $\boldsymbol{\delta\tilde{y}}(\omega)$

$$i\omega\boldsymbol{\delta\tilde{y}}(\omega) = \mathcal{G}\boldsymbol{\delta\tilde{s}}(\omega) + \mathcal{J}\boldsymbol{\delta\tilde{y}}(\omega) + \mathcal{B}\tilde{\boldsymbol{\xi}}(\omega), \tag{A3}$$

$$\boldsymbol{\delta\tilde{y}}(\omega) = (i\omega\mathbb{I}_n - \mathcal{J})^{-1}\left(\mathcal{G}\boldsymbol{\delta\tilde{s}}(\omega) + \mathcal{B}\tilde{\boldsymbol{\xi}}(\omega)\right). \tag{A4}$$

Using the convolution theorem to take the Fourier transform of Eq. A2, and comparing the result to Eq. A4, now shows that $\mathcal{F}\{e^{\mathcal{J}(t-t')}\} = (i\omega\mathbb{I}_n - \mathcal{J})^{-1}$. We obtain for the power-spectra of the network components

$$\begin{aligned}\mathcal{S}_y(\omega) &= \langle \delta\tilde{\boldsymbol{y}}(-\omega)\delta\tilde{\boldsymbol{y}}(\omega)^T\rangle,\\ &= \mathbb{G}(-\omega)\mathcal{S}_s(\omega)\mathbb{G}(\omega)^T + |\mathbb{N}(\omega)|^2,\end{aligned} \tag{A5}$$

with the matrices of frequency dependent gains $\mathbb{G}(\omega) \equiv (i\omega\mathbb{I}_n - \mathcal{J})^{-1}\mathcal{G}$, and frequency dependent noise $\mathbb{N}(\omega) \equiv (i\omega\mathbb{I}_n - \mathcal{J})^{-1}\mathcal{B}$. The cross terms vanish because the fluctuations of the external signal are uncorrelated from the internal noise. Furthermore, the power spectrum of a white noise process is constant, and all the noise terms are independent of one another, such that the spectral density of the noise vector is the identity matrix $\langle \tilde{\boldsymbol{\xi}}(-\omega)\tilde{\boldsymbol{\xi}}(\omega)^T\rangle = \mathbb{I}_m$. We also need to consider the cross-spectra between the signals and the network components, specifically we will need the spectra from the network to the signals

$$\begin{aligned}\mathcal{S}_{y\to s}(\omega) &= \langle \delta\tilde{\boldsymbol{y}}(-\omega)\delta\tilde{\boldsymbol{s}}(\omega)^T\rangle,\\ &= \mathbb{G}(-\omega)\mathcal{S}_s(\omega).\end{aligned} \tag{A6}$$

From Eq. A5 and Eq. A6 we can obtain all necessary correlation functions and (co-)variances, by taking the inverse Fourier transform of the component of interest (for a variance we can directly set $t = 0$). The advantage of using this form, is that the contribution of each signal and of the noise terms appear separately. When we are for example interested in a variance that is only caused by noise, we can omit the terms depending on the signal power spectra, and vice versa. Moreover, the power spectra are usually simpler in form than the corresponding correlation functions.

The covariance and auto-correlation matrices can also be found by solving Eq. A2 directly in the time domain; the solutions are shown here for completeness. For a derivation, see for example the work by Vennettilli et al. [5]. In this case it is most convenient to include the signals as system components, we thus have a new Jacobian $\mathcal{J}'$ and a new noise strength matrix $\mathcal{B}'$ which include all network components and the signals themselves. The covariance matrix $\mathcal{C}$ is then obtained by solving the Lyapunov equation

$$\mathcal{J}'\mathcal{C} + \mathcal{C}\mathcal{J}'^T + \mathcal{B}'\mathcal{B}'^T = \mathbf{0}, \tag{A7}$$

and the correlation matrix is given by

$$\mathcal{C}(\tau) = e^{\mathcal{J}'\tau}\mathcal{C} \qquad \text{for } \tau > 0. \tag{A8}$$

### Appendix B: Signals and statistics

#### 1. Markovian signal

For the Markovian ligand concentration dynamics we use a 1-dimensional OU-process

$$\delta\dot{\ell} = -\delta\ell/\tau_\ell + \eta_\ell(t), \tag{B1}$$

where the ligand concentration is defined in terms of the deviation from its mean $\delta\ell = \ell(t) - \bar{\ell}$. The correlation time is give by $\tau_\ell$, and the noise $\eta_\ell(t)$ is derived from a unit white noise process $\eta_\ell(t) \equiv \sigma_\ell\sqrt{2/\tau_\ell}\xi(t)$, such that $\langle\eta_\ell(t)\eta_\ell(t')\rangle = 2\sigma_\ell^2/\tau_\ell\delta(t-t')$. We obtain for the steady-state auto-correlation using Eq. A7 and Eq. A8:

$$\langle\delta\ell(\tau)\delta\ell(0)\rangle = \sigma_\ell^2 e^{-\tau/\tau_\ell}. \tag{B2}$$

#### 2. Non-Markovian signal

Not all ligand concentration trajectories encountered by cells are expected to be Markovian. For example, *E. coli* swims in its environment with a speed which exhibits persistence. This leads to an auto-correlation function for the concentrations' derivative which does not decay instantaneously [6]. To model such a persistent signal, we use the classical model of a particle in a harmonic well

$$\begin{aligned} \delta\dot{\ell} &= v(t), \\ \dot{v} &= -\omega_0^2\delta\ell(t) - v(t)/\tau_v + \eta_v(t), \end{aligned} \tag{B3}$$

where $\omega_0 = \sqrt{k/m}$, with $k$ the spring constant and $m$ the mass of the particle, $\tau_v$ is a relaxation timescale, and $\eta_v(t) = \sigma_v\sqrt{2/\tau_v}\xi(t)$, with $\xi(t)$, as used throughout, a Gaussian white noise process of unit variance, and $\sigma_v$ the standard deviation of $v$. If the signal would obey the fluctuation-dissipation relation, then $m\sigma_v^2 = k_\mathrm{B}T$, but since the biochemical signal could very well be generated via an active process this relation may not hold. This process can be expressed as a 2-dimensional OU-process with:

$$\mathcal{J} = \begin{pmatrix} 0 & 1 \\ -\omega_0^2 & -1/\tau_v \end{pmatrix}, \tag{B4}$$

$$\mathcal{B} = \begin{pmatrix} 0 & 0 \\ 0 & \sigma_v\sqrt{2/\tau_v} \end{pmatrix}. \tag{B5}$$

We find for the covariance matrix, using Eq. A7:

$$\mathcal{C} = \begin{pmatrix} \sigma_\ell^2 & \sigma_{\ell v} \\ \sigma_{\ell v} & \sigma_v^2 \end{pmatrix} = \sigma_v^2 \begin{pmatrix} 1/\omega_0^2 & 0 \\ 0 & 1 \end{pmatrix}. \tag{B6}$$

Using Eq. A8 we obtain the auto-correlation matrix in the overdamped regime, $\tau_v^{-1} > 2\omega_0$,

$$\mathcal{C}(\tau) = \begin{pmatrix} \langle \delta\ell(\tau)\delta\ell(0)\rangle & \langle \delta\ell(\tau)\delta v(0)\rangle \\ \langle \delta v(\tau)\delta\ell(0)\rangle & \langle \delta v(\tau)\delta v(0)\rangle \end{pmatrix},$$

$$= \begin{pmatrix} \sigma_\ell^2 e^{-\mu\tau/2}\left(\cosh(\rho\tau) + \frac{\mu}{2\rho}\sinh(\rho\tau)\right) & \sigma_v^2 e^{-\mu\tau/2}\frac{1}{\rho}\sinh(\rho\tau) \\ -\sigma_v^2 e^{-\mu\tau/2}\frac{1}{\rho}\sinh(\rho\tau) & \sigma_v^2 e^{-\mu\tau/2}\left(\cosh(\rho\tau) - \frac{\mu}{2\rho}\sinh(\rho\tau)\right) \end{pmatrix}, \tag{B7}$$

where $\rho = \sqrt{\mu^2/4 - \omega_0^2}$, with $\mu = \tau_v{}^{-1}$. The range of ligand concentrations which *E. coli* might encounter is very large, based on the dissociation constants of the inactive and active receptor conformations, which for the Tar-MeAsp receptor ligand combination respectively are $K_\mathrm{D}^\mathrm{I} = 18\mu\mathrm{M}$ and $K_\mathrm{D}^\mathrm{A} = 2900\mu\mathrm{M}$ [7, 8]. This suggests that the variance in the ligand concentration is very large relative to that of the derivative of the ligand concentration, which is set by the swimming behaviour of the cell. For this reason we specifically focus on the limit where $\omega_0 \to 0$, which corresponds to a vanishingly small spring constant, or a harmonic potential which becomes extremely wide. The variance in the concentration $\sigma_\ell^2$ then diverges, the normalized correlation functions in this limit are

$$\lim_{\omega_0\to 0}\begin{pmatrix} \frac{\langle \delta\ell(\tau)\delta\ell(0)\rangle}{\sigma_\ell^2} & \frac{\langle \delta\ell(\tau)\delta v(0)\rangle}{\sigma_\ell\sigma_v} \\ \frac{\langle \delta v(\tau)\delta\ell(0)\rangle}{\sigma_\ell\sigma_v} & \frac{\langle \delta v(\tau)\delta v(0)\rangle}{\sigma_v^2} \end{pmatrix} = \begin{pmatrix} 1 & 0 \\ 0 & e^{-\mu\tau} \end{pmatrix}. \tag{B8}$$

#### Appendix C: Information bottleneck framework and solutions

Anticipating future environmental conditions allows for timely adaptation. However, storing information costs resources such as proteins, energy and time, and not all information in the past ligand concentrations will be relevant for predicting the signal's future state. Assuming that resources are in limited supply, this means that cells must be efficient in which, and how much information they store. This is elegantly captured in the Information Bottleneck Method (IBM), which describes the problem of maximizing the information on the future signal while minimizing the information on the past signal that is stored in the network output, from which the future input is predicted [9]. The objective function for the prediction of a variable of interest $z_\tau \equiv z(t+\tau)$ is:

$$\max_{P(X_0|\boldsymbol{L}_p)} : \quad \mathcal{L} = I(x_0; z_\tau) - \gamma I(x_0; \boldsymbol{L}_p). \tag{C1}$$

The value of the sensing system output at the current time $t$ is $x_0 \equiv x(t)$. The variable of interest $z_\tau$ at a future time $t+\tau$ is the future concentration $\ell_\tau \equiv \ell(t+\tau)$ for the Markovian signal, and the future concentration derivative $v_\tau \equiv v(t+\tau)$ for the non-Markovian signal. Since the system of interest needs to predict one signal characteristic (either the future signal value or its derivative), one output component is sufficient for encoding the required information, as we describe in more detail below. The vector $\boldsymbol{L}_p = (\delta\ell(0), \delta\ell(-\Delta t), \ldots, \delta\ell(-(N-1)\Delta t))^T$ is the past trajectory of ligand concentrations of length $N$, discretized with timestep $\Delta t$. The mutual information between the current system output and the future property of interest is the predictive information $I_\mathrm{pred} \equiv I(x_0; z_\tau)$, and the mutual information between the current system output and the past ligand concentration trajectory is the past information $I_\mathrm{past} \equiv I(x_0; \boldsymbol{L}_p)$. The Lagrange multiplier $\gamma$ sets the relative cost of storing past information over obtaining predictive information. Given a value of $\gamma$, Eq. C1 is maximized by optimizing the mapping of the past ligand concentration trajectory $\boldsymbol{L}_p$ onto the current output $x_0$. Since, by the data processing inequality, we have $I_\mathrm{past} \geq I_\mathrm{pred}$, for $\gamma = 1$ the objective function is maximized by $I_\mathrm{past} = I_\mathrm{pred} = 0$. As $\gamma$ is decreased both the past and predictive information increase, and the parametric curve in the $I_\mathrm{past} - I_\mathrm{pred}$ plane that arises is the information bound. For $\gamma = 0$ there is no cost to storing past information. The predictive information is then only limited by the amount of information contained in the past about the future signal property: $I_\mathrm{pred} \leq I(\boldsymbol{L}_p; z_\tau)$.

##### 1. Gaussian information bottleneck

In general equation C1 can be difficult to solve, as all mappings from $\boldsymbol{L}_p$ to $X_0$ are allowed. However, the problem becomes analytically tractable when the joint probability distribution of $\boldsymbol{L}_p$ and $z_\tau$ is a multivariate Gaussian. Here, we follow the procedure of Chechik and coworkers to obtain this mapping [10]. In the Gaussian model, the optimal mapping from $\boldsymbol{L}_p$ to $x_0$ is a linear one [10]

$$x_0 = \boldsymbol{A}\boldsymbol{L}_p + \xi; \quad \xi \sim N(0, \sigma_\xi^2), \tag{C2}$$

where $\boldsymbol{A}$ is a row vector which determines how strongly each entry in $\boldsymbol{L}_p$ contributes to the scalar output $X_0$ at any point in time. The random variable $\xi$ is the noise added to the signal due to the stochastic nature of the mapping; it is a Gaussian random variable independent of $\boldsymbol{L}_p$ with 0 mean and variance $\sigma_\xi^2$. Finding the optimal mapping from $\boldsymbol{L}_p$ to $x_0$ corresponds to finding the optimal combination of $\boldsymbol{A}$ and $\sigma_\xi^2$. It can be shown that for any pair $(\boldsymbol{A}, \sigma_\xi^2)$, there exists a pair $(\boldsymbol{A}', 1)$ which yields the same values for $I_{\text{past}}$ and $I_{\text{pred}}$ after maximization of Eq. C1 [10]. Therefore, we can set $\sigma_\xi^2 = 1$ without altering the information curve.

To obtain the information bound, we rewrite Eq. C1 using the definition of the mutual information between Gaussian random variables:

$$\mathcal{L} = \frac{1}{2}\log(\sigma_x^2/\sigma_{x|z}^2) - \gamma\frac{1}{2}\log(\sigma_x^2/\sigma_{x|L}^2), \tag{C3}$$

with the total variance $\sigma_x^2$ in the output $x_0$, the output variance conditional on the future signal property $\sigma_{x|z}^2 \equiv \sigma_{x|z_\tau}^2$, and the output variance conditional on the complete history of ligand concentrations $\sigma_{x|L}^2 \equiv \sigma_{x|\boldsymbol{L}_p}^2$. The latter is just the variance caused by the intrinsic noise, $\sigma_{x|L}^2 = \sigma_\xi^2 = 1$. The total variance in $x_0$ can be expressed in terms of the mapping vector $\boldsymbol{A}$ and the variance in the past signal using Eq. C2, $\sigma_x^2 = \boldsymbol{A}\boldsymbol{\Sigma}_L\boldsymbol{A}^T + 1$, where $\boldsymbol{\Sigma}_L \equiv \boldsymbol{\Sigma}_{\boldsymbol{L}_p}$ is the covariance matrix of the past ligand concentration trajectory $\boldsymbol{L}_p$. To express the output variance conditional on the future signal property $z_\tau$ we use the Schur complement formula, which in general form reads:

$$\boldsymbol{\Sigma}_{x|y} = \boldsymbol{\Sigma}_x - \boldsymbol{\Sigma}_{xy}\boldsymbol{\Sigma}_y^{-1}\boldsymbol{\Sigma}_{yx}, \tag{C4}$$

where $\boldsymbol{\Sigma}_{yx} = \boldsymbol{\Sigma}_{xy}^T$. Using this formula to rewrite $\sigma_{x|z}^2$, and then using the linear relation from Eq. C2 again, we obtain $\sigma_{x|z}^2 = \boldsymbol{A}\boldsymbol{\Sigma}_{L|z}\boldsymbol{A}^T + 1$.

Filling in the expressions for the variances in $\mathcal{L}$ (Eq. C3) gives:

$$\mathcal{L} = \frac{1}{2}\left((1-\gamma)\log\left(\left|\boldsymbol{A}\boldsymbol{\Sigma}_L\boldsymbol{A}^T + 1\right|\right) - \log\left(\left|\boldsymbol{A}\boldsymbol{\Sigma}_{L|z}\boldsymbol{A}^T + 1\right|\right)\right). \tag{C5}$$

For any symmetric matrix $\boldsymbol{C}$ we have $\frac{\delta}{\delta A}\log|\boldsymbol{A}\boldsymbol{C}\boldsymbol{A}^T| = \left(\boldsymbol{A}\boldsymbol{C}\boldsymbol{A}^T\right)^{-1} 2\boldsymbol{A}\boldsymbol{C}$, such that we obtain for the derivative of $\mathcal{L}$ to $\boldsymbol{A}$:

$$\frac{\delta\mathcal{L}}{\delta\boldsymbol{A}} = (1-\gamma)\frac{\boldsymbol{A}\boldsymbol{\Sigma}_L}{\boldsymbol{A}\boldsymbol{\Sigma}_L\boldsymbol{A}^T + 1} - \frac{\boldsymbol{A}\boldsymbol{\Sigma}_{L|z}}{\boldsymbol{A}\boldsymbol{\Sigma}_{L|z}\boldsymbol{A}^T + 1}. \tag{C6}$$

In our case $\boldsymbol{A}$ is a row vector, and both denominators are thus scalars. We find the maximum of $\mathcal{L}$ by equating its derivative to 0, which gives:

$$\boldsymbol{A}\boldsymbol{\Sigma}_{L|z}\boldsymbol{\Sigma}_L^{-1} = (1-\gamma)\frac{\boldsymbol{A}\boldsymbol{\Sigma}_{L|z}\boldsymbol{A}^T + 1}{\boldsymbol{A}\boldsymbol{\Sigma}_L\boldsymbol{A}^T + 1}\boldsymbol{A}. \tag{C7}$$

For this equality to hold $\boldsymbol{A}$ must either be identically 0, or a left eigenvector of the matrix $\boldsymbol{\Sigma}_{L|z}\boldsymbol{\Sigma}_L^{-1}$ with eigenvalue:

$$\lambda = (1-\gamma)\frac{\boldsymbol{A}\boldsymbol{\Sigma}_{L|z}\boldsymbol{A}^T + 1}{\boldsymbol{A}\boldsymbol{\Sigma}_L\boldsymbol{A}^T + 1}. \tag{C8}$$

Here, we note that if the signal statistics is sufficiently rich and the prediction complexity sufficiently large (because, for example, multiple signal characteristics need to be predicted), then the matrix $\boldsymbol{\Sigma}_{L|z}\boldsymbol{\Sigma}_L^{-1}$ has multiple eigenvectors with non-trivial eigenvalues $0 < \lambda_i < 1$ [10]. This reflects the idea that storing the past information that is necessary to enable this complex prediction task may require multiple output components, i.e. an output vector $\boldsymbol{x}$, where each output component has an integration kernel given by one of the eigenvectors of $\boldsymbol{\Sigma}_{L|z}\boldsymbol{\Sigma}_L^{-1}$ [10]. However, for Markovian signals only one eigenvector with non-trivial eigenvalue $0 < \lambda < 1$ emerges, which means that one output component is sufficient to encode the required information. For the non-Markovian signals studied here, $\boldsymbol{\Sigma}_{L|z}\boldsymbol{\Sigma}_L^{-1}$ has two eigenvectors if both the future value and its derivative need to be predicted (and $z = (\ell_\tau, v_\tau)$); to optimally predict both features from the current output, two output components are then required, provided $I_{\text{past}}$ is sufficiently large. However, here we consider the scenario that only the future derivative needs to be predicted, in which case only one non-trivial eigenvector emerges, and one output component is sufficient for encoding the required information. We leave the problem of predicting multiple signal features via multiple output components for future work.

We can define the optimal mapping $\boldsymbol{A} = ||A||\boldsymbol{\nu}$ where $\boldsymbol{\nu}$ is the normalized left eigenvector of $\boldsymbol{\Sigma}_{L|z}\boldsymbol{\Sigma}_L^{-1}$ corresponding to its smallest eigenvalue, $0 < \lambda < 1$. The magnitude can be found by solving Eq. C8 for $||A||$, using from Eq. C7 that $\lambda\boldsymbol{\nu}\boldsymbol{\Sigma}_L\boldsymbol{\nu}^T = \boldsymbol{\nu}\boldsymbol{\Sigma}_{L|z}\boldsymbol{\nu}^T$. This gives for the optimal mapping:

$$\boldsymbol{A}^{\text{opt}} = \begin{cases} \sqrt{\frac{1-\gamma-\lambda}{\boldsymbol{\nu}_1\boldsymbol{\Sigma}_L\boldsymbol{\nu}_1^T\lambda\gamma}}\boldsymbol{\nu}_1 & \text{for} \quad 0 < \lambda < 1-\gamma, \\ 0 & \text{for} \quad 1-\gamma \leq \lambda \leq 1. \end{cases} \tag{C9}$$

We can substitute $||A||^2 = (1-\gamma-\lambda)/(\boldsymbol{\nu}\boldsymbol{\Sigma}_L\boldsymbol{\nu}^T\lambda\gamma)$ in the definitions for the mutual information to express them in terms of $\lambda$ and $\gamma$. For the past information we obtain:

$$\begin{aligned} I_{\text{past}} &= \frac{1}{2}\log\left(||A||^2\boldsymbol{\nu}\boldsymbol{\Sigma}_L\boldsymbol{\nu}^T + 1\right), \\ &= \frac{1}{2}\log\left(\frac{1-\gamma}{\gamma}\frac{1-\lambda}{\lambda}\right). \end{aligned} \tag{C10}$$

And for the predictive information:

$$\begin{aligned} I_{\text{pred}} &= \frac{1}{2}\log\left(||A||^2\boldsymbol{\nu}\boldsymbol{\Sigma}_L\boldsymbol{\nu}^T + 1\right) - \frac{1}{2}\log\left(||A||^2\boldsymbol{\nu}\boldsymbol{\Sigma}_{L|\ell_\tau}\boldsymbol{\nu}^T + 1\right), \\ &= I_{\text{past}} - \frac{1}{2}\log\left(\frac{1-\lambda}{\gamma}\right), \\ &= \frac{1}{2}\log\left(\frac{1-\gamma}{\lambda}\right). \end{aligned} \tag{C11}$$

#### 2. Markovian signal

To obtain the information bound for prediction of the future ligand concentration of a Markovian signal, we need to determine the eigenvalues and vectors of the matrix (see Eqs. C7 and C8)

$$\boldsymbol{W} = \boldsymbol{\Sigma}_{L|\ell_\tau}\boldsymbol{\Sigma}_L^{-1}. \tag{C12}$$

Using the Schur complement formula (Eq. C4) to rewrite the conditional matrix gives $\boldsymbol{\Sigma}_{L|\ell_\tau} = \boldsymbol{\Sigma}_L - \boldsymbol{\Sigma}_{L\ell_\tau}\boldsymbol{\Sigma}_{L\ell_\tau}^T/\sigma_\ell^2$. Then defining the normalized matrices $\boldsymbol{R}_L = \boldsymbol{\Sigma}_L/\sigma_\ell^2$ and $\boldsymbol{R}_{L\ell_\tau} = \boldsymbol{\Sigma}_{L\ell_\tau}/\sigma_\ell^2$ we find

$$\boldsymbol{W} = \mathbb{I}_N - \boldsymbol{R}_{L\ell_\tau}\boldsymbol{R}_{L\ell_\tau}^T\boldsymbol{R}_L^{-1}. \tag{C13}$$

where $N$ is the length of the input trajectory $\boldsymbol{L}_p$. The correlation matrix of the past trajectory is symmetric with entries $\boldsymbol{R}_L^{(i,j)} = \exp(-|i-j|\Delta t/\tau_\ell)$, where $\Delta t$ is the discretization timestep of the past trajectory $\boldsymbol{L}_p$ and $i$ ranges from 1 to $N$. This is a Kac-Murdock-Szegö matrix, and its inverse is known:

$$\boldsymbol{R}_L^{-1} = \frac{1}{1-e^{2\Delta t/\tau_\ell}}\begin{pmatrix} 1 & -e^{-\Delta t/\tau_\ell} & 0 & \dots & \dots & 0 \\ -e^{-\Delta t/\tau_\ell} & 1+e^{-2\Delta t/\tau_\ell} & -e^{-\Delta t/\tau_\ell} & \dots & \dots & 0 \\ 0 & -e^{-\Delta t/\tau_\ell} & 1+e^{-2\Delta t/\tau_\ell} & \ddots & \dots & 0 \\ \vdots & \vdots & \ddots & \ddots & \ddots & \vdots \\ 0 & \dots & \dots & -e^{-\Delta t/\tau_\ell} & 1+e^{-2\Delta t/\tau_\ell} & -e^{-\Delta t/\tau_\ell} \\ 0 & \dots & \dots & 0 & -e^{-\Delta t/\tau_\ell} & 1 \end{pmatrix}. \tag{C14}$$

Note that the inverse matrix is tridiagonal. The length $N$ cross-correlation vector between past trajectory and future concentration has entries $\boldsymbol{R}_{L\ell_\tau}^{(i)} = \exp(-(\tau+(i-1)\Delta t)/\tau_\ell)$. The product of the correlation matrices is surprisingly simple:

$$\boldsymbol{R}_{L\ell_\tau}\boldsymbol{R}_{L\ell_\tau}^T\boldsymbol{R}_L^{-1} = e^{-2\tau/\tau_\ell}\begin{pmatrix} 1 & 0 & \dots & 0 \\ e^{-\Delta t/\tau_\ell} & 0 & \dots & 0 \\ \vdots & \vdots & \ddots & \vdots \\ e^{-(N-1)\Delta t/\tau_\ell} & 0 & \dots & 0 \end{pmatrix}. \tag{C15}$$

Using this result we can straightforwardly determine the eigenvalues,

$$|\boldsymbol{W} - \lambda \mathbb{I}_N| = 0,$$

$$\left| \begin{pmatrix} 1-\lambda - e^{-2\tau/\tau_\ell} & 0 & \dots & 0 \\ -e^{-(\tau+\Delta t)/\tau_\ell} & 1-\lambda & \dots & 0 \\ \vdots & \vdots & \ddots & \vdots \\ -e^{-(\tau+(N-1)\Delta t)/\tau_\ell} & 0 & \dots & 1-\lambda \end{pmatrix} \right| = 0. \tag{C16}$$

The only contribution to the determinant comes from the diagonal, and the only nontrivial eigenvalue is thus $\lambda = 1 - e^{-2\tau/\tau_l}$. The optimal mapping is thus onto a one-dimensional scalar output $x_0$. The corresponding left eigenvector is given by

$$\boldsymbol{\nu}_1 \boldsymbol{W} = (1 - e^{-2\tau/\tau_l})\boldsymbol{\nu}_1, \tag{C17}$$

which holds for $\boldsymbol{\nu}_1 = \begin{pmatrix} 1 & 0 & \dots & 0 \end{pmatrix}$. The optimal mapping for the prediction of a one-dimensional OU-process is thus to copy its most recent value. This agrees with intuition as for any Markovian process, all the information about the future signal is contained in the most recent value. For a continuous input signal (rather than a discretized signal), and a continuous integration kernel $k(t)$ (rather than a mapping vector $\boldsymbol{A}$), this means that the optimal integration kernel is $k^{\text{opt}}(t) = a\delta(t)$.

#### 3. Non-Markovian signal

To find the optimal mapping for the prediction of the derivative of a non-Markovian signal, based on its history of ligand concentrations, we need to find the eigenvalues and vectors of the matrix

$$\begin{aligned} \boldsymbol{W} &= \boldsymbol{\Sigma}_{L|v_\tau} \boldsymbol{\Sigma}_L^{-1}, \\ &= \mathbb{I}_N - \frac{1}{\sigma_v^2} \boldsymbol{\Sigma}_{Lv_\tau} \boldsymbol{\Sigma}_{Lv_\tau}^T \boldsymbol{\Sigma}_L^{-1}. \end{aligned} \tag{C18}$$

The covariance matrix of the past trajectory is symmetric with entries $\boldsymbol{\Sigma}_L^{(i,j)} = \langle \delta\ell(0)\delta\ell(|i-j|\Delta t)\rangle$ where both $i$ and $j$ range from 1 to $N$, the past trajectory length. The covariance vector between past trajectory and future derivative has entries $\boldsymbol{\Sigma}_{Lv_\tau}^{(i,j)} = \langle \delta\ell(0)\delta v(\tau + (i-1)\Delta t)\rangle$. Both the concentration auto-correlation function, and the concentration to future derivative cross-correlation function, are shown in Eq. B7.

To better understand the optimal mapping of this signal we numerically investigate the eigenvalues of the matrix $\boldsymbol{W}$. For the prediction of $v_\tau$, there is only one non-trivial eigenvalue. Like for the Markovian signal, this shows that for the prediction of the derivative of this non-Markovian signal, the optimal mapping is always onto a scalar output. The non-trivial eigenvalue $\lambda$ decreases with the discretization timestep $\Delta t$ and is minimal for $\Delta t \to 0$ Fig. 1. In this limit, $\lambda$ has the same magnitude for any $N \geq 2$, see Fig. 1. A smaller eigenvalue $\lambda$ corresponds to larger past and predictive information and a larger ratio $I_{\text{pred}}/I_{\text{past}}$ (Eq. C10 and Eq. C11), given any value of the Lagrange multiplier $\gamma$. For the optimal mapping we must thus have $N \geq 2$ and $\Delta t \to 0$, where $N$ sets both the past trajectory and the mapping vector length. Because increasing the length above two does not yield an improvement in the value of $\lambda_1$ we focus on $N = 2$.

The fact that to reach the optimum we must have $N = 2$ and $\Delta t \to 0$, shows that the optimal kernel $A$ takes an instantaneous measurement of a combination of the most recent ligand concentration, and its derivative. This can be understood as follows, for a trajectory of length two, the mapping vector also has length two, $\boldsymbol{A} = ||A||(\hat{w}_1, \hat{w}_2)$, with $\sqrt{\hat{w}_1^2 + \hat{w}_2^2} = 1$. We can then express the linear mapping of $\boldsymbol{L}_p$ to $x_0$ (Eq. C2) as:

$$x_0 = ||A|| \left[ (\hat{w}_1 + \hat{w}_2)\delta\ell(0) - \hat{w}_2 \Delta t \frac{\delta\ell(0) - \delta\ell(-\Delta t)}{\Delta t} \right] + \xi, \tag{C19}$$

This expression shows that, as $\Delta t \to 0$, the two entries of $\boldsymbol{A}$ combine both the most recent signal value and the most recent derivative to generate $x_0$. This is intuitive because the signal is completely defined by its concentration and derivative (Eq. B3). For this reason, and to obtain analytical insight into the optimal weights, we inspect the final two entries of the past ligand concentration trajectory in the limit $\Delta t \to 0$, which defines the past signal in terms of its most recent concentration and derivative

$$\boldsymbol{S}_0 \equiv \begin{pmatrix} \delta\ell(0) & v(0) \end{pmatrix}^T. \tag{C20}$$

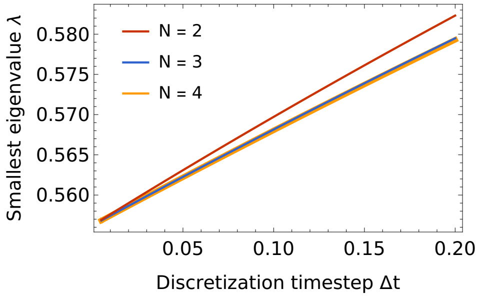


FIG. 1. **The smallest eigenvalue of the IB matrix is minimal for $N \geq 2$ and $\Delta t \to 0$.** A smaller eigenvalue corresponds to a larger ratio $I_{\text{pred}}/I_{\text{past}}$ for any given value of the Lagrange multiplier $\gamma$. Parameters: friction timescale $\tau_v^{-1} = 0.862 s^{-1}$ as determined in [6], prediction interval $\tau = \tau_v$, and $\omega_0 = 0.4 s^{-1}$ such that the system is slightly overdamped.

Because the signal is Markovian in the joint properties $\delta\ell$ and $v$, the vector $\boldsymbol{S}_0$ contains the same information as the trajectory $\boldsymbol{L}_p$. The past information is now the mutual information between $x_0$ and $\boldsymbol{S}_0$, i.e. $I_{\text{past}} = I(x_0; \boldsymbol{S}_0)$. The output $x_0$ can then also be written as a projection of $\boldsymbol{S}_0$ via the alternative mapping vector $\tilde{\boldsymbol{A}} = ||A||(\hat{a}, \hat{b})$:

$$x_0 = ||A|| \left(\hat{a}\delta\ell(0) + \hat{b}v(0)\right) + \xi. \tag{C21}$$

Comparison with Eq. C19 shows how the components of $\tilde{\boldsymbol{A}}$ relate back to those in $\boldsymbol{A}$,

$$\hat{w}_1 = \hat{a} + \hat{b}/\Delta t, \tag{C22}$$

$$\hat{w}_2 = -\hat{b}/\Delta t. \tag{C23}$$

To obtain the optimal mapping vector $\tilde{\boldsymbol{A}}$ the matrix of signal statistics of which the eigenvalues and -vectors need to be determined is

$$\boldsymbol{W} = \boldsymbol{\Sigma}_{\boldsymbol{s}|v_\tau} \boldsymbol{\Sigma}_{\boldsymbol{s}}^{-1}, \tag{C24}$$

with

$$\boldsymbol{\Sigma}_{\boldsymbol{s}} = \begin{pmatrix} \sigma_\ell^2 & 0 \\ 0 & \sigma_v^2 \end{pmatrix}, \tag{C25}$$

$$\boldsymbol{\Sigma}_{\boldsymbol{s}|v_\tau} = \boldsymbol{\Sigma}_{\boldsymbol{s}} - \frac{1}{\sigma_v^2} \boldsymbol{\Sigma}_{\boldsymbol{s}v_\tau} \boldsymbol{\Sigma}_{\boldsymbol{s}v_\tau}^T, \tag{C26}$$

$$\boldsymbol{\Sigma}_{\boldsymbol{s}v_\tau} = \begin{pmatrix} \langle \delta\ell(0)\delta v(\tau)\rangle \\ \langle \delta v(0)\delta v(\tau)\rangle \end{pmatrix}. \tag{C27}$$

We thus obtain

$$\boldsymbol{W} = \mathbb{I} - \begin{pmatrix} \frac{\langle \delta\ell(0)\delta v(\tau)\rangle^2}{\sigma_\ell^2 \sigma_v^2} & \frac{\langle \delta\ell(0)\delta v(\tau)\rangle \langle \delta v(0)\delta v(\tau)\rangle}{\sigma_v^4} \\ \frac{\langle \delta\ell(0)\delta v(\tau)\rangle \langle \delta v(0)\delta v(\tau)\rangle}{\sigma_\ell^2 \sigma_v^2} & \frac{\langle \delta v(0)\delta v(\tau)\rangle^2}{\sigma_v^4} \end{pmatrix}. \tag{C28}$$

This matrix has one nontrivial eigenvalue, $\lambda = 1 - \frac{\langle \delta v(0)\delta v(\tau)\rangle^2}{\sigma_v^4} - \frac{\langle \delta\ell(0)\delta v(\tau)\rangle^2}{\sigma_\ell^2\sigma_v^2}$, which depends on the normalized correlation functions between on the one hand the current concentration or derivative, and on the other hand the future derivative. The corresponding left eigenvector is

$$\boldsymbol{\nu}_1 = Q^{-1} \begin{pmatrix} \frac{1}{\sigma_\ell} \frac{\langle \delta\ell(0)\delta v(\tau)\rangle}{\sigma_\ell \sigma_v} & \frac{1}{\sigma_v} \frac{\langle \delta v(0)\delta v(\tau)\rangle}{\sigma_v^2} \end{pmatrix}, \tag{C29}$$

where $Q$ normalizes the vector.Using the linear mapping $x_0 = ||A|| \boldsymbol{\nu}_1 \boldsymbol{S}_0 + \xi$, and defining $G \equiv ||A||/Q$, shows that the optimal output should be generated as follows

$$x_0^{\text{opt}} = G \left( \frac{\langle \delta\ell(0)\delta v(\tau)\rangle}{\sigma_\ell \sigma_v} \frac{\delta\ell(0)}{\sigma_\ell} + \frac{\langle \delta v(0)\delta v(\tau)\rangle}{\sigma_v^2} \frac{v(0)}{\sigma_v} \right) + \xi. \tag{C30}$$

Clearly, the optimal mapping depends on the (normalized) cross-correlation coefficient $\rho_{\ell_0 v_\tau} \equiv \langle \delta\ell(0)\delta v(\tau)\rangle/(\sigma_\ell \sigma_v)$ between the current concentration $\delta\ell(0)$ and future derivative $\delta v(\tau)$, and the cross-correlation coefficient $\rho_{v_0 v_\tau}$ between the current derivative $\delta v(0)$ and future derivative $\delta v(\tau)$. Indeed, to optimally predict the future derivative, the cell should also use the current concentration and not only its current derivative. However, in the limit that the range of concentrations sensed becomes very large, corresponding to $\omega_0 \to 0$, the current concentration is no longer correlated with the future derivative, and $\rho_{\ell_0 v_\tau} \to 0$ (Eq. B8). In this limit, $\hat{a} = 0$ and $\hat{b} = 1$, and the kernel becomes a perfectly adaptive, derivative-taking kernel:

$$\lim_{\omega_0 \to 0} x_0^{\text{opt}} = ||A|| v(0) + \xi. \tag{C31}$$

If we translate this back to the vector $||A||(\hat{w}_1, \hat{w}_2)$, operating on a ligand concentration trajectory $\boldsymbol{L}_p$, the optimal weights become $\hat{w}_1 = -\hat{w}_2$.

### Appendix D: Past and predictive information for linear signalling networks

In order to address how close biochemical networks can come to the information bounds derived above, we here describe how we obtain the past and predictive information for any linear (biochemical) network. We then use the resulting general expressions to compute the past and predictive information for the push-pull network and the chemotaxis system of the main text.

For any linear network the output can be written as

$$\delta x(t) = \int_{-\infty}^{t} ds\, k(t-s)\delta\ell(s) + \eta_x(t). \tag{D1}$$

The mapping kernel $k(t)$ is a property of the network and describes how the input signal is mapped onto the output. The noise term $\eta_x(t)$ is a sum of convolutions over all white noise processes in the network and corresponding network mapping functions, see Eq. A2. The variance in the output can generally be split up in a part caused by the signal and a part cause by the noise, and we have

$$\begin{aligned} \sigma_x^2 &= \int_{-\infty}^{t} ds \int_{-\infty}^{t} ds' k(t-s)k(t-s')\langle \delta\ell(s)\delta\ell(s')\rangle + \sigma_{\eta_x}^2, \\ &= \sigma_{x|\eta}^2 + \sigma_{x|L}^2, \end{aligned} \tag{D2}$$

where $\sigma_{x|\eta}^2$ is the signal variance, i.e. all noise terms are fixed, and $\sigma_{x|L}^2$ is the noise variance, i.e. the complete history of the signal is fixed. Using this decomposition we find for the past information, which is the mutual information between the current output and the complete signal history,

$$I_{\text{past}}(x_0; \boldsymbol{L}_p) = \frac{1}{2}\log\left(\frac{\sigma_x^2}{\sigma_{x|L}^2}\right) = \frac{1}{2}\log(1+\text{SNR}), \tag{D3}$$

where the signal-to-noise ratio is defined as $\text{SNR} = \sigma_{x|\eta}^2/\sigma_{x|L}^2$. Using the same definition for the mutual information when deriving the predictive information between current output and future ligand concentration, we obtain

$$\begin{aligned} I_{\text{pred}}(x_0; \ell_\tau) &= \frac{1}{2}\log\left(\frac{\sigma_x^2}{\sigma_{x|\ell_\tau}^2}\right), \\ &= \frac{1}{2}\log\left(1 + \frac{\sigma_{x|\eta}^2}{\sigma_{x|L}^2}\right) - \frac{1}{2}\log\left(1 + \frac{\sigma_{x|\eta}^2 - \langle \delta x(0)\delta\ell(\tau)\rangle^2/\sigma_\ell^2}{\sigma_{x|L}^2}\right), \\ &= I_{\text{past}} - \frac{1}{2}\log(1+\text{cSNR}). \end{aligned} \tag{D4}$$

In the second line we used the Schur complement formula, Eq. C4, to decompose the variance in the output conditioned on the future signal: $\sigma_{x|\ell_\tau}^2 = \sigma_x^2 - \langle \delta x(0)\delta\ell(\tau)\rangle^2/\sigma_\ell^2$. The quantity $\sigma_{x|\eta}^2 - \langle \delta x(0)\delta\ell(\tau)\rangle^2/\sigma_\ell^2$ can be understood as follows: the first term $\sigma_{x|\eta}^2$ is the contribution to the total variance of the output $\sigma_x^2$ that comes from the signal variations, while the second term quantifies the variance in the output that is correlated with the future input. The difference is thus the variance in the output coming from the signal variations that are not correlated with the future input. The ratio in the second logarithm can thus be understood as a conditional SNR that quantifies the part of the signal to noise ratio that does *not* contain information about the future signal. This becomes more clear when considering its form in terms of the mapping kernel and signal correlation functions. For any linear signalling network we have

$$\sigma^2_{x|\eta} - \langle \delta x(0)\delta\ell(\tau)\rangle^2/\sigma^2_\ell = \int_{-\infty}^0 ds \int_{-\infty}^0 ds' k(-s)k(-s') \left( \langle \delta\ell(s)\delta\ell(s')\rangle - \frac{\langle \delta\ell(\tau)\delta\ell(s')\rangle\langle\delta\ell(\tau)\delta\ell(s)\rangle}{\sigma^2_\ell} \right), \tag{D5}$$

where the term in parentheses is the conditional variance in the past signal trajectory given a future value, $\mathbf{\Sigma}_{L|\ell_\tau}$. The form in Eq. D4 thus tells us that the predictive information is equal to the past information, minus the bits that do not contain information about the future ligand concentration. This difference is indeed the part of the past information that does contain predictive information about the future signal.

Although the expression above (Eq. D4) nicely relates the past and predictive information, a more straightforward way of obtaining the predictive information is by expressing it directly in terms of the correlation between the current output and the future ligand concentration:

$$I_{\rm pred}(x_0;\ell_\tau) = \frac{1}{2}\log\left(\frac{\sigma^2_x}{\sigma^2_{x|\ell_\tau}}\right) = -\frac{1}{2}\log\left(1 - \frac{\langle \delta x(0)\delta\ell(\tau)\rangle^2}{\sigma^2_x\sigma^2_\ell}\right), \tag{D6}$$

where we again used the Schur complement formula to rewrite $\sigma^2_{x|\ell_\tau}$. Written this way we thus see that the predictive information depends on the normalized correlation between the current network output and the future ligand concentration. We can simply exchange the future ligand concentration for the future derivative when considering the chemotaxis network.

To compute the past information for linear signalling networks we use Eq. D3, and we thus need to compute the SNR. To compute the predictive information for the prediction of a future ligand concentration, we need to compute the 'future correlation function' $\langle \delta x(0)\delta\ell(\tau)\rangle$. For the prediction of the future derivative we need $\langle \delta x(0)\delta v(\tau)\rangle$.

### Appendix E: Push-pull network

We consider a push-pull network that consists of a phosphorylation-dephosphorylation cycle downstream of a receptor. When bound to ligand, the receptor itself or its associated kinase, such as CheA in *E. coli*, catalyzes the phosphorylation of a readout protein $x$, like CheY. Active readout molecules $x^*$ can decay spontaneously or be deactivated by an enzyme (phosphatase), such as CheZ in *E. coli.* This cycle is driven by the turnover of fuel such as ATP. We recognize that inside the living cell, the chemical driving is typically large: for example, the free energy of ATP hydrolysis is about $20k_{\rm B}T$, which means that the system essentially operates in the irreversible regime [11, 12]. This system consists of the following reactions:

$$\mathrm{R} + \mathrm{L} \underset{\mathrm{k}_-}{\overset{\mathrm{k}_+}{\rightleftharpoons}} \mathrm{RL} \tag{E1}$$

$$\mathrm{RL} + \mathrm{x} \xrightarrow{\mathrm{k_f}} \mathrm{RL} + \mathrm{x}^* \tag{E2}$$

$$\mathrm{X}^* \xrightarrow{\mathrm{k_r}} \mathrm{X} \tag{E3}$$

Both the total number of receptors $R_{\rm T} = R + RL$ and read-out molecules $X_{\rm T} = X + X^*$ are conserved moieties. The chemical Langevin equations of this system are:

$$\dot{RL} = [R_T - RL(t)]\ell(t)k_+ - RL(t)k_- + B_c(RL,\ell)\xi_c(t), \tag{E4}$$

$$\dot{x}^* = [X_T - x^*(t)]RL(t)k_{\rm f} - x^*(t)k_{\rm r} + B_x(RL,x^*)\xi_x(t), \tag{E5}$$

where $RL$ is the number of bound receptors, $x^*$ the number of phosphorylated read-out molecules, and $\xi_i$ denote independent Gaussian white noise with unit variance, $\langle \xi_i(t)\xi_j(t')\rangle = \delta_{ij}\delta(t-t')$. The noise strengths are $B_c(RL,\ell) = \sqrt{(R_T - RL(t))\ell(t)k_+ + RL(t)k_-}$ and $B_x(RL,x^*) = \sqrt{(X_T - x^*(t))RL(t)k_{\rm f} + x^*(t)k_{\rm r}}$. The steady-state fraction of ligand-bound receptors is $p \equiv \overline{RL}/R_T = \bar{\ell}/(\bar{\ell}+K_{\rm D})$ with the dissociation constant $K_{\rm D} = k_-/k_+$, and the steady-state fraction of phosphorylated readout molecules is $f \equiv \bar{x}^*/X_{\rm T} = pR_{\rm T}/(pR_{\rm T} + k_{\rm r}/k_{\rm f})$.

In the linear-noise approximation, expanding Eqs. E4 and E5 to first order around their steady state, the equations become

$$\delta\dot{RL} = b\,\delta\ell(t) - \delta RL(t)/\tau_{\rm c} + \eta_c(t), \tag{E6}$$

$$\delta\dot{x}^* = \gamma\,\delta RL(t) - \delta x^*(t)/\tau_{\rm r} + \eta_x(t). \tag{E7}$$

The parameters $b = R_\mathrm{T}p(1-p)/(\bar{\ell}\tau_\mathrm{c})$ and $\gamma = X_\mathrm{T}f(1-f)/(R_\mathrm{T}p\tau_\mathrm{r})$ are effective rates of receptor-ligand binding and readout phosphorylation, respectively. The decay rate of correlations in the receptor-ligand binding state is ${\tau_\mathrm{c}}^{-1} = \bar{\ell}k_+ + k_-$, and that of the readout phosphorylation state is ${\tau_\mathrm{r}}^{-1} = pR_\mathrm{T}k_\mathrm{f} + k_\mathrm{r}$. The rescaled white noise processes have strengths $\langle\eta_c^2\rangle = B_c^2 = 2R_\mathrm{T}p(1-p)/\tau_\mathrm{c}$ and $\langle\eta_x^2\rangle = B_x^2 = 2X_\mathrm{T}f(1-f)/\tau_\mathrm{r}$.

#### 1. Model statistics

The relevant quantity to compute the past information is the variance in the output, decomposed into the part caused by signal variation and the part caused by noise. To compute the predictive information we further need the correlation function between the current output and a future ligand concentration $\langle\delta\ell(\tau)\delta x^*(0)\rangle$. These quantities can be obtained via their Fourier transforms, as in Eq. A5 and Eq. A6. The matrices describing the properties of the signalling network are, as defined below Eq. A1,

$$\mathcal{G} = \begin{pmatrix} b \\ 0 \end{pmatrix}, \tag{E8}$$

$$\mathcal{J} = \begin{pmatrix} -{\tau_\mathrm{c}}^{-1} & 0 \\ \gamma & -{\tau_\mathrm{r}}^{-1} \end{pmatrix}, \tag{E9}$$

$$\mathcal{B} = \begin{pmatrix} \sqrt{\langle\eta_c^2\rangle} & 0 \\ 0 & \sqrt{\langle\eta_x^2\rangle} \end{pmatrix} = \begin{pmatrix} \sqrt{2R_\mathrm{T}p(1-p)/\tau_\mathrm{c}} & 0 \\ 0 & \sqrt{2X_\mathrm{T}f(1-f)/\tau_\mathrm{r}} \end{pmatrix}. \tag{E10}$$

A useful property of the network is the matrix exponential of its Jacobian, which in Fourier space is (see Eq. A2 and Eq. A4)

$$\begin{aligned} \mathcal{F}\{e^{\mathcal{J}t}\}(\omega) &= (i\omega\mathbb{I}_2 - \mathcal{J})^{-1}, \\ &= \begin{pmatrix} \frac{1}{1/\tau_\mathrm{c}+i\omega} & 0 \\ \frac{\gamma}{(1/\tau_\mathrm{c}+i\omega)(1/\tau_\mathrm{r}+i\omega)} & \frac{1}{1/\tau_\mathrm{r}+i\omega} \end{pmatrix}. \end{aligned} \tag{E11}$$

We then have $\mathbb{G}(\omega) = \mathcal{F}\{e^{\mathcal{J}t}\}(\omega)\mathcal{G}$ and $\mathbb{N}(\omega) = \mathcal{F}\{e^{\mathcal{J}t}\}(\omega)\mathcal{B}$, see also Eq. A4 and Eq. A5. The integration kernel that maps the ligand concentration onto the output of the push-pull network, see Eq. D1, is given by the inverse Fourier transform of the second entry of $\mathbb{G}(\omega)$, which is the frequency dependent gain, $\tilde{g}_{\ell\to x}(\omega)$, from $\ell$ to $x$:

$$\begin{aligned} k(t) \equiv \mathcal{F}^{-1}\{\tilde{g}_{\ell\to x}(\omega)\} &= b\gamma\tau_\mathrm{c}\tau_\mathrm{r}\frac{1}{\tau_\mathrm{r}-\tau_\mathrm{c}}\left(e^{-t/\tau_\mathrm{r}} - e^{-t/\tau_\mathrm{c}}\right), \\ &= X_\mathrm{T}f(1-f)(1-p)/\bar{\ell}\frac{1}{\tau_\mathrm{r}-\tau_\mathrm{c}}\left(e^{-t/\tau_\mathrm{r}} - e^{-t/\tau_\mathrm{c}}\right), \end{aligned} \tag{E12}$$

The so-called static gain of the network is the integral of this kernel over all time, $\bar{g}_{\ell\to x} \equiv \int_0^\infty k(t)dt = X_\mathrm{T}f(1-f)(1-p)/\bar{\ell}$. This parameter quantifies how much a step change in the input concentration changes the steady-state level of the output: $\bar{g}_{\ell\to x} = \partial\bar{x}^*/\partial\bar{\ell}$. We will use this parameter in the statistical quantities that follow. The static gain is also given by $\bar{g}_{\ell\to x} = \bar{g}_{\ell\to RL}\bar{g}_{RL\to x}$, with $\bar{g}_{\ell\to RL} = p(1-p)R_\mathrm{T}/\bar{\ell}$ the static gain from $\bar{\ell}$ to $RL$ and $\bar{g}_{RL\to x} = f(1-f)X_\mathrm{T}/(pR_\mathrm{T})$ the static gain from $RL$ to $x^*$.

We model the Markovian ligand concentration as a 1-dimensional OU process Eq. B1, which has the following power spectrum

$$S_\ell(\omega) = \langle|\delta\ell(\omega)|^2\rangle = \frac{2\sigma_\ell^2/\tau_\ell}{1/\tau_\ell^2+\omega^2}. \tag{E13}$$

This yields the following expression for the power spectra (see Eq. A5):

$$\mathbb{G}(-\omega)S_\ell(\omega)\mathbb{G}(\omega)^T = b^2\begin{pmatrix} \frac{1}{1/{\tau_\mathrm{c}}^2+\omega^2} & \gamma\frac{1}{1/\tau_\mathrm{r}-i\omega}\frac{1}{1/{\tau_\mathrm{c}}^2+\omega^2} \\ \gamma\frac{1}{1/\tau_\mathrm{r}+i\omega}\frac{1}{1/{\tau_\mathrm{c}}^2+\omega^2} & \gamma^2\frac{1}{1/{\tau_\mathrm{r}}^2+\omega^2}\frac{1}{1/{\tau_\mathrm{c}}^2+\omega^2} \end{pmatrix}\frac{2\sigma_\ell^2/\tau_\ell}{1/\tau_\ell^2+\omega^2} \tag{E14}$$

$$|\mathbb{N}(\omega)|^2 = \langle\eta_c^2\rangle\begin{pmatrix} \frac{1}{1/{\tau_\mathrm{c}}^2+\omega^2} & \gamma\frac{1}{1/\tau_\mathrm{r}-i\omega}\frac{1}{1/{\tau_\mathrm{c}}^2+\omega^2} \\ \gamma\frac{1}{1/\tau_\mathrm{r}+i\omega}\frac{1}{1/{\tau_\mathrm{c}}^2+\omega^2} & \gamma^2\frac{1}{1/{\tau_\mathrm{r}}^2+\omega^2}\frac{1}{1/{\tau_\mathrm{c}}^2+\omega^2} + \frac{\langle\eta_x^2\rangle}{\langle\eta_c^2\rangle}\frac{1}{1/{\tau_\mathrm{r}}^2+\omega^2} \end{pmatrix} \tag{E15}$$

We thus have for the power spectrum of the read-out:

$$S_x(\omega) = \tilde{g}^2_{\ell\to x}(\omega)S_\ell(\omega) + N_x^2(\omega)$$
$$= \frac{2b^2\gamma^2\sigma_\ell^2/\tau_\ell}{(1/\tau_\mathrm{r}{}^2+\omega^2)(1/\tau_\mathrm{c}{}^2+\omega^2)(1/\tau_\ell^2+\omega^2)} + \frac{\gamma^2\langle\eta_c^2\rangle}{(1/\tau_\mathrm{r}{}^2+\omega^2)(1/\tau_\mathrm{c}{}^2+\omega^2)} + \frac{\langle\eta_x^2\rangle}{1/\tau_\mathrm{r}{}^2+\omega^2}, \tag{E16}$$

The variance in the read-out $\sigma_x^2 = 1/(2\pi)\int_{-\infty}^{\infty} S_x(\omega)$ is hence given by

$$\sigma_x^2 = \sigma^2_{x|\eta} + \sigma^2_{x|L}$$
$$= \bar{g}^2_{\ell\to x}\frac{1+\tau_\mathrm{r}/\tau_\ell+\tau_\mathrm{r}/\tau_\mathrm{c}}{(1+\tau_\mathrm{c}/\tau_\ell)(1+\tau_\mathrm{r}/\tau_\ell)(1+\tau_\mathrm{r}/\tau_\mathrm{c})}\sigma_\ell^2 + \bar{g}^2_{RL\to x}R_\mathrm{T}p(1-p)\frac{1}{1+\tau_\mathrm{r}/\tau_\mathrm{c}} + X_\mathrm{T}f(1-f),$$
$$= \underbrace{\bar{g}^2_{\ell\to x}\frac{1+\tau_\mathrm{r}/\tau_\ell+\tau_\mathrm{r}/\tau_\mathrm{c}}{(1+\tau_\mathrm{c}/\tau_\ell)(1+\tau_\mathrm{r}/\tau_\ell)(1+\tau_\mathrm{r}/\tau_\mathrm{c})}}_{\text{dynamical gain}}\sigma_\ell^2 + X_\mathrm{T}f(1-f)\left(1+\bar{g}_{\ell\to x}\frac{\bar{\ell}}{R_\mathrm{T}p}\frac{1}{1+\tau_\mathrm{r}/\tau_\mathrm{c}}\right), \tag{E17}$$

where $\bar{g}_{RL\to x} = \gamma\tau_r = X_\mathrm{T}f(1-f)/(R_\mathrm{T}p)$ is the static gain from the receptor to the readout. The expression above gives insight into the role of the different network components in shaping the noise in the readout. It can be seen that the contribution from the signal variance $\sigma_\ell^2$ to $\sigma_x^2$ is determined by the static gain $\bar{g}^2_{\ell\to x}$, which is proportional to $X_\mathrm{T}$, and a factor that only depends on ratios of timescales. Their product is the dynamical gain, which decreases monotonically with $\tau_\mathrm{r}$. The intrinsic noise in the phosphorylation state of the read-outs leads to the noise term $X_\mathrm{T}f(1-f)$, which cannot be averaged out. The noise arising from ligand binding and unbinding increases with the static gain, but can be mitigated by increasing the number of receptors or the integration time $\tau_\mathrm{r}$. The latter strategy is what we call time-averaging.

The signal-to-noise ratio $\mathrm{SNR} = \sigma^2_{x|\eta}/\sigma^2_{x|L}$ can straightforwardly be obtained from Eq. E17. This is the quantity that sets the magnitude of the past information, see Eq. D3. To determine the predictive information we need to compute the correlation function from the current output to the future ligand concentration $\langle\delta x(0)\delta\ell(\tau)\rangle$. This requires the cross-spectrum from output to ligand concentration, which is given by (Eq. A6)

$$\tilde{g}_{\ell\to x}(-\omega)S_\ell(\omega) = \frac{b\gamma}{(1/\tau_\mathrm{c}-i\omega)(1/\tau_\mathrm{r}-i\omega)}\frac{2\sigma_\ell^2/\tau_\ell}{1/\tau_\ell^2+\omega^2}. \tag{E18}$$

From this power spectrum we obtain the required correlation function by taking the inverse Fourier transfrom:

$$\langle\delta x(0)\delta\ell(\tau)\rangle = \mathcal{F}^{-1}\{\tilde{g}_{\ell\to x}(-\omega)S_\ell(\omega)\},$$
$$= \frac{\bar{g}_{\ell\to x}\sigma_\ell^2}{(1+\tau_\mathrm{c}/\tau_\ell)(1+\tau_\mathrm{r}/\tau_\ell)}e^{-\tau/\tau_\ell}. \tag{E19}$$

This correlation function thus decays exponentially with the prediction interval $\tau$ at a rate $\tau_\ell^{-1}$, just as the signal auto-correlation. The (squared) correlation coefficient, which sets $I_\mathrm{pred}$, is given by $\langle\delta x(0)\delta\ell(\tau)\rangle^2/(\sigma_\ell^2\sigma_x^2) = \rho^2_{\ell x}e^{-2\tau/\tau_\ell}$, with the (squared) instantaneous correlation coefficient (for convenience given as its inverse)

$$\rho_{\ell x}^{-2} = \frac{\bar{\ell}^2}{\sigma_\ell^2}\left(1+\frac{\tau_\mathrm{r}}{\tau_\ell}\right)^2\left(1+\frac{\tau_\mathrm{c}}{\tau_\ell}\right)^2\left(\frac{1}{X_Tf(1-f)(1-p)^2} + \frac{1}{R_Tp(1-p)(1+\tau_\mathrm{r}/\tau_\mathrm{c})} + \frac{\sigma_\ell^2}{\bar{\ell}^2}\frac{1+\tau_\mathrm{r}/\tau_\ell+\tau_\mathrm{r}/\tau_\mathrm{c}}{(1+\tau_\mathrm{c}/\tau_\ell)(1+\tau_\mathrm{r}/\tau_\ell)(1+\tau_\mathrm{r}/\tau_\mathrm{c})}\right). \tag{E20}$$

When the right-hand-side is minimized, the correlation is thus maximized. This expression shows that increasing $X_\mathrm{T}$ and $R_\mathrm{T}$ always increases the instantaneous correlation coefficient, and that the fraction of phosphorylated readout molecules in steady state that maximizes the correlation coefficient is $f = 1/2$.

#### 2. Past and predictive information of the push-pull network

Using the quantitites computed above, we can determine both the past and the predictive information. For the past information we use Eq. D3, whith the SNR from Eq. E17:

$$\mathrm{SNR} = \sigma^2_{x|\eta}/\sigma^2_{x|L} = (1-p)\frac{\sigma_\ell^2}{\bar{\ell}^2}\frac{1+\tau_\mathrm{r}/\tau_\ell+\tau_\mathrm{r}/\tau_\mathrm{c}}{(1+\tau_\mathrm{c}/\tau_\ell)(1+\tau_\mathrm{r}/\tau_\ell)(1+\tau_\mathrm{r}/\tau_\mathrm{c})}\Bigg/\left(\frac{1}{X_\mathrm{T}f(1-f)(1-p)} + \frac{1}{R_\mathrm{T}p(1+\tau_\mathrm{r}/\tau_\mathrm{c})}\right).$$

The predictive information is a function of the correlation between the current output and the future ligand concentration, Eq. D6. This correlation can be decomposed into the instantaneous correlation coefficient and an exponential decay on the timescale of the ligand concentration fluctuations, Eq. E19. We thus obtain for the predictive information,

$$I_{\rm pred}(x_0;\ell_\tau) = -\frac{1}{2}\log(1-\rho^2_{\ell x}e^{-2\tau/\tau_\ell}). \tag{E21}$$

The instantaneous correlation coefficient $\rho^2_{\ell x}$ is given in Eq. E20. From Eq. E21 it also becomes clear that while the value of the predictive information depends on the forecast interval $\tau$, the optimal design of the network that maximizes the predictive information, determined by the optimal ratio $X_{\rm T}/R_{\rm T}$, the optimal integration time $\tau_{\rm r}$, and the optimal ligand-bound receptor fraction $p$, does not depend on the forecast interval $\tau$.

#### 3. Optimal resource allocation

Increasing the number of receptor or readout molecules always increases the precision with which the cell can predict a signal (see Eq. E20). However, when the total resource pool is constrained, the cell has to choose whether it makes more receptors or more readout molecules. To find the optimal ratio of read-out to receptor molecules we, can use the $C = AR_{\rm T} + BX_{\rm T}$ to express $X_{\rm T}$ and $R_{\rm T}$ in terms of the total cost $C$ and the ratio $X_{\rm T}/R_{\rm T}$:

$$X_{\rm T} = C\frac{X_{\rm T}/R_{\rm T}}{A+BX_{\rm T}/R_{\rm T}}, \tag{E22}$$

$$R_{\rm T} = C\frac{1}{A+BX_{\rm T}/R_{\rm T}}. \tag{E23}$$

The factors $A$ and $B$ set the cost of receptors and readout molecules, respectively. Substituting these expressions for $X_{\rm T}$ and $R_{\rm T}$ into the expression for the correlation coefficient between the output and ligand concentration (Eq. E20), setting the derivative of the resulting expression with respect to $X_{\rm T}/R_{\rm T}$ to zero, and solving for $X_{\rm T}/R_{\rm T}$ gives

$$\begin{aligned}(X_{\rm T}/R_{\rm T})^{\rm opt} &= \sqrt{\left(1+\frac{\tau_{\rm r}}{\tau_{\rm c}}\right)\frac{p}{1-p}\frac{1}{f(1-f)}\frac{A}{B}},\\ &= 2\sqrt{p/(1-p)}\sqrt{1+\tau_{\rm r}/\tau_{\rm c}},\end{aligned} \tag{E24}$$

where for the second line we used $A = B = 1$ and $f = f^{\rm opt} = 1/2$. This is the optimal ratio of readout to receptor molecules in the push-pull network, given an integration time $\tau_{\rm r}$ and a steady state fraction of ligand-bound receptors $p$. Perhaps surprisingly, this optimal ratio $(X_{\rm T}/R_{\rm T})^{\rm opt}$ maximizes, for a given $\tau_{\rm r}$ and $p$, not only the predictive information, but also the past information. This is because the ratio $X_{\rm T}/R_{\rm T}$ determines, together with $\tau_{\rm r}$ and $p$, the interval $\Delta$ for sampling the ligand-binding state of the receptor: when the ratio $X_{\rm T}/R_{\rm T}$ obeys Eq. E24, the readout molecules sample each receptor molecule roughly once every correlation time: $\Delta \sim \tau_{\rm c}$ [11, 12]. Eq. E24 is thus a statement about optimally extracting the information that is encoded in the receptor-ligand binding history, both concerning the past information and the predictive information. This is illustrated in Fig. 2.

#### 4. Operating costs diverge when approaching the information bound

The precision of any sensing device is limited by the resources that are devoted to it. The cost function we consider in this work is

$$C = \lambda(R_{\rm T}+X_{\rm T}) + c_1X_{\rm T}\Delta\mu/\tau_{\rm r}. \tag{E25}$$

The first term is the maintenance cost; this is the cost of producing new network components at the growth rate $\lambda$. The second term is the operating cost and describes the chemical power that is necessary to run the network; it depends on the flux through the network, $X_{\rm T}/\tau_{\rm r}$, and the free-energy drop $\Delta\mu$ over a full cycle of phosphorylation and dephosphorylation, given by the free energy of ATP hydrolysis. The coefficient $c_1$ describes the relative energetic cost of synthesising the components during the cell cycle, versus that of running the system. In the main text we consider the case where $c_1 \to 0$. Here we will investigate how close cells can come to the information bound when $c_1$ is finite, thus including the chemical power cost of running the network.

It is clear from *Eq.* $E25$ that for finite $c_1$ the operating cost diverges when $\tau_{\rm r} \to 0$. Because the optimal IBM solutions are instantaneous, this is precisely the limit in which the network must be to reach the information bound.

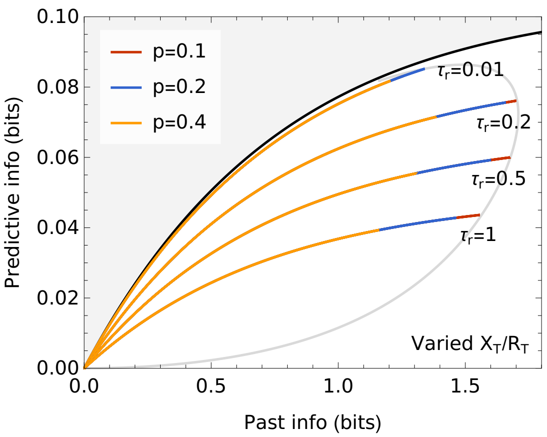


FIG. 2. **The past and predictive information are maximized by the same ratio $X_{\rm T}/R_{\rm T}$ and fraction $p$.** The information plane, showing the information bound in black, and the isocost line $C = 10^4$ in gray. To construct the coloured lines in this figure the ratio $X_{\rm T}/R_{\rm T}$ has been varied from zero to a value beyond the optimal value that maximizes $I_{\rm past}$ and $I_{\rm pred}$. This is done for several values of the receptor occupancy $p$ ($p = 0.1$ in red, $p = 0.2$ in blue, $p = 0.4$ in orange), and for several values of $\tau_{\rm r}$ (indicated in the figure). When $X_{\rm T}/R_{\rm T}$ reaches its optimal value, both $I_{\rm past}$ and $I_{\rm pred}$ are maximal. When the ratio is increased further the system moves back to the origin via the same coordinates. Only the integration time $\tau_{\rm r}$ meaningfully distinuishes between strategies that maximize predictive or past information, or that approach the information bound. The reason is that $X_{\rm T}/R_{\rm T}$, together with $\tau_{\rm r}$ and $p$, control the optimal extraction of information that is encoded in the receptor-ligand binding history, both concerning $I_{\rm past}$ and $I_{\rm pred}$. The gray isocost line is obtained by varying $\tau_{\rm r}$, while maximizing for each $\tau_{\rm r}$ the correlation coefficient given by Eq. E20; the latter is done by substituting Eq. E24 into Eq. E20 and numerically optimizing the resulting expression over $p$. The isocost line gives the region of $I_{\rm past}$ and $I_{\rm pred}$ that is accessible for a given resource cost $C$. Parameter values are $A = B = 1$, $f = 1/2$, $(\sigma_\ell/\bar{\ell})^2 = 10^{-2}$, $\tau_{\rm c}/\tau_\ell = 10^{-2}$.

As a consequence, when we consider the operating costs, the push-pull network can only be at the information bound when $(I_{\rm past}, I_{\rm pred}) \to (0, 0)$ or $C \to \infty$ (Fig. 3A). The system can mitigate the operating costs by decreasing $X_{\rm T}$, because this decreases the flux through the cycle. However, this also decreases the gain and thus, eventually, any information transduced through the network. In the limit that both $X_{\rm T}$ and $\tau_{\rm r}$ approach zero, the system approaches the information bound at the origin, see both Fig. 3A and B. More generally, when the running costs are taken into account, the system time averages more (i.e., $\tau_{\rm r}$ rises), because frequent measurements are now even more costly. Still, $\tau_{\rm r}$ decreases as the total resource availability $C$ grows.

### Appendix F: Chemotaxis network

The evidence is mounting that in the *E. coli* chemotaxis system, receptors cooperatively control the activity of the kinase CheA [13–16]. Furthermore, the kinase activity is adaptive due to the methylation of inactive receptors [17, 18]. A widely used approach to describe the effects of receptor cooperativity and methylation on kinase activity, has been to employ the Monod-Wyman-Changeux (MWC) model [6–8, 15, 19–22]. We will follow this approach and, more specifically, model the chemotaxis system as described by Tu and colleagues [23]. In this model, each receptor can switch between an active and inactive conformational state. Moreover, receptors are partitioned into clusters of equal size $N$. In the spirit of the MWC model, receptors within a cluster switch conformation in concert, so that each cluster is either active or inactive [19]. Furthermore, it is assumed that receptor-ligand binding and conformational switching are faster than the other timescales in the system. The probability for the kinase, i.e. the receptor cluster, to be active, is then described by:

$$a(\ell, m) = \frac{1}{1 + \exp(\Delta F_T(\ell, m))}, \tag{F1}$$

where $\Delta F_T(\ell, m)$ is the total free-energy difference between the active and inactive state, which is a function of the ligand concentration $\ell(t)$ and the methylation level of the cluster $m(t)$. The simplest model adopted here assumes a linear dependence of the total free-energy difference on the free-energy difference arising from ligand binding and methylation:

$$\Delta F_T(\ell, m) = -\Delta E_0 + N(\Delta F_\ell(\ell) + \Delta F_m(m)), \tag{F2}$$

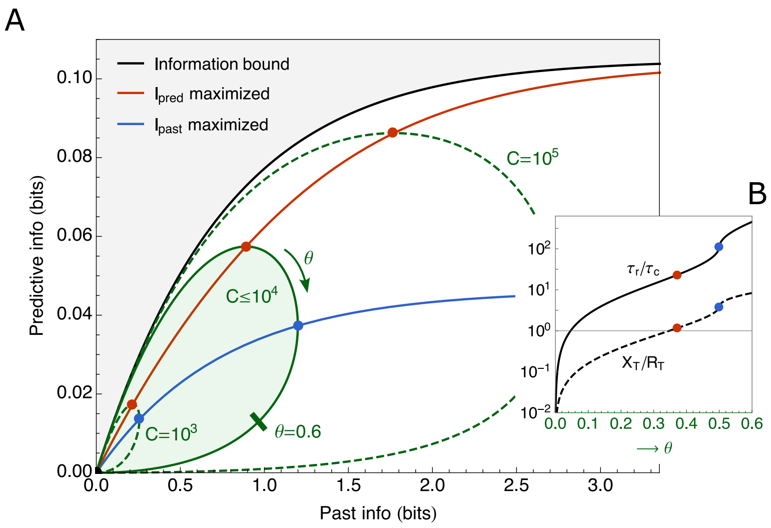


FIG. 3. **Due to diverging operating costs the push-pull network only reaches the information bound for infinite resource availability.** (A) In green, the region of accessible predictive and past information in the push-pull network under a resource constraint $C = \lambda(R_T + X_T) + c_1 X_T \Delta\mu/\tau_r$, with $\lambda = 1$ and $c_1 = 1/\Delta\mu$, corresponding to a cell doubling time of roughly 20min [11]. The black line is the information bound; the red and blue dots mark the points where $I_{\rm pred}$ and $I_{\rm past}$ are maximized, respectively, under a resource constraint $C$; the red and blue lines connect these points, respectively, for increasing $C$. The accesible region for $C \leq 10^4$ and the isocost lines for $C = 10^3$ and $C = 10^5$ have been obtained as described under Fig. 2. The forecast interval has been set to one signal correlation time in the future: $\tau = \tau_\ell$. (B) The integration time over the receptor correlation time, $\tau_r/\tau_c$, and the ratio of the number of readout and receptor molecules, $X_T/R_T$, as a function of the distance $\theta$ along the iscocost line for $C = 10^4$ in panel A. For $\theta \to 0$, both $\tau_r$ and $X_T$ go to zero, thus reducing both $I_{\rm past}$ and $I_{\rm pred}$ to zero. Other parameter values in both panels are $f = f^{\rm opt} = 1/2$, $(\sigma_\ell/\bar{\ell})^2 = 10^{-2}$, $\tau_c/\tau_\ell = 10^{-2}$.

where the free-energy difference due to ligand binding is

$$\Delta F_\ell(\ell) = \ln(1 + \ell(t)/K_D^I) - \ln(1 + \ell(t)/K_D^A). \tag{F3}$$

Between the two states the cluster has an altered dissociation constant, which is denoted $K_D^I$ for the inactive state, and $K_D^A$ for the active state. The free-energy difference due to methylation has been experimentally shown to depend approximately linearly on the methylation level [15]:

$$\Delta F_m(m) = \tilde{\alpha}(\bar{m} - m(t)). \tag{F4}$$

We assume that inactive receptors are irreversibly methylated, and active receptors irreversibly demethylated, with zero-order ultrasensitive kinetics [23–25]. The dynamics of the methylation level of the $i^{\rm th}$ receptor cluster is then given by:

$$\dot{m}_i = (1 - a_i(\ell, m_i))k_R - a_i(\ell, m_i)k_B + B_{m_i}(a_i)\xi(t), \tag{F5}$$

with $B_m^{(i)}(a_i) = \sqrt{(1 - a_i(\ell, m_i))k_R + a_i(\ell, m_i)k_B}$, and unit white noise $\xi(t)$. These dynamics indeed give rise to perfect adaptation, since from this equation we find that the steady state cluster activity is given by $p \equiv \bar{a} = 1/(1 + k_B/k_R)$, thus indeed independent of the ligand concentration.

Finally, active receptors catalyze phosphorylation of read-out molecules, and phosphorylated read-out molecules decay at a constant rate. We have

$$\dot{x}^* = \sum_{i=1}^{R_T} a_i(t)(X_T - x^*(t))k_f - x^*(t)k_r + B_x(a_i, x^*)\xi(t), \tag{F6}$$

where $R_T$ is the total number of receptor *clusters*. The steady state fraction of phosphorylated read-outs is given by $f \equiv \bar{x}^*/X_T = (1 + k_r/(k_f R_T p))^{-1}$.

#### 1. Linear dynamics

We again do a first order approximation around the steady state, defining all variables in terms of deviations from their mean: $\delta\ell(t) = \ell(t) - \bar{\ell}$, $\delta m(t) = m(t) - \bar{m}$ and $\delta a(t) = a(t) - p$. The linear form of this model has previously been studied in for example [23] and [25]. We obtain for the linear dynamics of the $i^{\text{th}}$ cluster activity

$$\delta a_i(t) = \alpha \delta m_i(t) - \beta \delta\ell(t), \tag{F7}$$

with $\alpha = \tilde{\alpha} N p(1-p)$ and $\beta = \kappa N p(1-p)$, with $\kappa = (\bar{\ell} + K_{\text{D}}^{\text{I}})^{-1} - (\bar{\ell} + K_{\text{D}}^{\text{A}})^{-1}$. For the methylation on the $i^{\text{th}}$ cluster and for the readout dynamics we then obtain, as a function of $\delta a(t)$,

$$\dot{\delta m_i} = -\delta a_i(t)/(\alpha\tau_{\text{m}}) + \eta_{m_i}(t), \tag{F8}$$

$$\dot{\delta x^*} = \gamma \sum_{i=1}^{R_{\text{T}}} \delta a_i(t) - \delta x^*(t)/\tau_r + \eta_x(t), \tag{F9}$$

where we have introduced the relaxation times $\tau_{\text{m}} = (\alpha(k_R + k_B))^{-1}$ for methylation and $\tau_{\text{r}} = (R_{\text{T}} p k_{\text{f}} + k_{\text{r}})^{-1}$ for phosphorylation. We have further defined the rate at which an active cluster phosphorylates the readout CheY: $\gamma = X_{\text{T}} f(1-f)/(p R_{\text{T}} \tau_{\text{r}})$. Substituting the expression for $\delta a_i$ in Eq. F7 into Eqs. F8 and F9, and expressing the dynamics in terms of the methylation on all clusters gives

$$\frac{d}{dt}\left(\sum_{i=1}^{R_T} \delta m_i\right) = -\sum_{i=1}^{R_T} \delta m_i/\tau_{\text{m}} + q\delta\ell(t)/(\alpha\tau_{\text{m}}) + \eta_m(t), \tag{F10}$$

$$\dot{\delta x^*} = -\delta x^*(t)/\tau_{\text{r}} - \gamma q \delta\ell(t) + \gamma\alpha \sum_{i=1}^{R_{\text{T}}} \delta m_i(t) + \eta_x(t), \tag{F11}$$

with $q = R_{\text{T}}\beta$ (see Eq. F7 for $\beta$). The rescaled white noise $\eta_m$ is the sum of the methylation noise on all receptor clusters, $\langle \eta_m^2 \rangle = 2R_{\text{T}} p(1-p)/(\alpha\tau_{\text{m}})$, where we have assumed that the methylation noise on the respective receptor clusters is independent. The phosphorylation noise has strength $\langle \eta_x^2 \rangle = 2X_{\text{T}} f(1-f)/\tau_{\text{r}}$.

#### 2. Parameter values

A large body of work has studied the parameters of the MWC model for the *E. coli* chemotaxis system. We have listed the parameters relevant for our model in table I. We choose the background concentration $\bar{\ell}$ to be in between $K_{\text{D}}^{\text{I}}$ and $K_{\text{D}}^{\text{A}}$, at $\bar{\ell} = 100\mu$M.

In this work we analyze the impact of the methylation timescale $\tau_{\text{m}}$, and the numbers of receptor clusters and readout molecules $R_{\text{T}}$ and $X_{\text{T}}$, on the past and predictive information. We therefore do not set them to a fixed value, but experimental estimates are listed in table II.

#### 3. Model statistics

Again we take the power spectrum route to determine the variance in the network output, the SNR, and the correlation coefficient between current output and the future signal. We consider the system to sense the non-Markovian ligand concentration defined in equation Eq. B3. Such a signal is characterized by both its concentration

TABLE I. **Measured *E. coli* chemotaxis parameter values.**

| *Parameter* | *Value* | *Source* | *Description* |
|---|---|---|---|
| $K_{\text{D}}^{\text{I}}$ | $18\mu$M | [7, 8] | MeAsp-Tar dissociation constant inactive receptor |
| $K_{\text{D}}^{\text{A}}$ | $2900\mu$M | [7, 8] | MeAsp-Tar dissociation constant active receptor |
| $N$ | $\sim 6$ | [7, 8, 15, 26] | Number of receptors per cluster |
| $\tilde{\alpha}$ | $2k_{\text{B}}T$ | [15] | Free energy change per added methyl group |
| $p$ | $\frac{1}{3}, \frac{1}{2}$ | [8, 15] | Steady state activity at $22°C$, $32°C$ |
| $\tau_{\text{r}}$ | $\sim 0.1s$ | [6, 11, 26] | Phosphorylation timescale |

TABLE II. **Approximate *E. coli* chemotaxis timescales and abundances.**

| *Parameter* | *Value* | *Source* | *Description* |
|---|---|---|---|
| $\tau_{\rm m}$ | $\sim 10s$ | [6, 15, 17] | Adaptation time |
| Tsr+Tar | $14000, 3300$ | [27] | Rich medium; RP437, OW1 strain |
| Tsr+Tar | $24000, 37000$ | [27] | Minimal medium; RP437, OW1 strain |
| CheY | $8200, 1400$ | [27] | Rich medium; RP437, OW1 strain |
| CheY | $6300, 14000$ | [27] | Minimal medium; RP437, OW1 strain |

and derivative, and the (cross-)power spectra of these properties are

$$\mathcal{S}_s(\omega) = \begin{pmatrix} S_\ell(\omega) & S_{\ell\to v}(\omega) \\ S_{v\to\ell}(\omega) & S_v(\omega) \end{pmatrix} = \begin{pmatrix} S_\ell(\omega) & i\omega S_\ell(\omega) \\ -i\omega S_\ell(\omega) & \omega^2 S_\ell(\omega) \end{pmatrix}, \tag{F12}$$

with

$$S_\ell(\omega) = \frac{2\sigma_v^2/\tau_v}{(\omega^2 + ((2\tau_v)^{-1}+\rho)^2)(\omega^2 + ((2\tau_v)^{-1}-\rho)^2)}, \tag{F13}$$

where $\rho = \sqrt{(4\tau_v^2)^{-1} - \omega_0^2}$. The chemotaxis signalling network is fully determined by the following matrices (Eq. A1)

$$\mathcal{G} = q \begin{pmatrix} 1/(\alpha\tau_{\rm m}) & 0 \\ -\gamma & 0 \end{pmatrix}, \tag{F14}$$

$$\mathcal{J} = \begin{pmatrix} -1/\tau_{\rm m} & 0 \\ \alpha\gamma & -1/\tau_{\rm r} \end{pmatrix}, \tag{F15}$$

$$\mathcal{B} = \begin{pmatrix} \sqrt{\langle\eta_m^2\rangle} & 0 \\ 0 & \sqrt{\langle\eta_x^2\rangle} \end{pmatrix}. \tag{F16}$$

The Fourier transform of the matrix exponential of the Jacobian is

$$\begin{aligned} \mathcal{F}\{e^{\mathcal{J}t}\} &= (i\omega\mathbb{I}_n - \mathcal{J})^{-1} \\ &= \begin{pmatrix} \frac{1}{1/\tau_{\rm m}+i\omega} & 0 \\ \frac{\alpha\gamma}{(1/\tau_{\rm m}+i\omega)(1/\tau_{\rm r}+i\omega)} & \frac{1}{1/\tau_{\rm r}+i\omega} \end{pmatrix}, \end{aligned} \tag{F17}$$

which allows us to determine the gain matrix via $\mathbb{G}(\omega) = \mathcal{F}\{e^{\mathcal{J}t}\}(\omega)\mathcal{G}$, and the noise matrix using $\mathbb{N}(\omega) = \mathcal{F}\{e^{\mathcal{J}t}\}(\omega)\mathcal{B}$; see also Eq. A4 and Eq. A5.

To gain more insight in the way in which the network maps the signal onto its output, we first study the integration kernels of the system. The integration kernel from ligand concentration to output is given by the inverse Fourier transform of element $(1,2)$ of the gain matrix $\mathbb{G}(\omega)$, which is

$$k(t) \equiv \mathcal{F}^{-1}\{\tilde{g}_{\ell\to x}(\omega)\} = \kappa N f(1-f)(1-p)X_{\rm T}\frac{1}{1-\tau_{\rm r}/\tau_{\rm m}}\left(\frac{1}{\tau_{\rm m}}e^{-\tau/\tau_{\rm m}} - \frac{1}{\tau_{\rm r}}e^{-\tau/\tau_{\rm r}}\right), \tag{F18}$$

with $\kappa = (\bar{\ell}+K_{\rm D}^{\rm I})^{-1} - (\bar{\ell}+K_{\rm D}^{\rm A})^{-1}$. Due to the adaptive nature of the network, the static gain from ligand concentration to output is zero: $\bar{g}_{\ell\to x} = \int_0^\infty k(t)dt = 0$; the long-time response to a step change in a constant input is zero. The kernel does indeed not change the output based on the input concentration directly, but instead takes a (time-averaged) derivative of the input (Fig. 4A). It is therefore useful to consider the kernel that maps the signal derivative onto the output. This kernel can be found by rearranging the expression for the output of a linear signalling network, Eq. D1. Disregarding the noise terms and integrating by parts gives

$$\int_{-\infty}^0 k(-t)\ell(t)dt = K(-t)\ell(t)|_{-\infty}^0 - \int_{-\infty}^0 K(-t)v(t)dt, \tag{F19}$$

where $v(t) \equiv \dot{\ell}$ and $K(t)$ is the primitive of $k(t)$. To make progress we first determine $K(t)$,

$$K(t) = \kappa N f(1-f)(1-p) X_T \frac{1}{1-\tau_{\rm r}/\tau_{\rm m}} \left(-e^{-\tau/\tau_{\rm m}} + e^{-\tau/\tau_{\rm r}}\right). \tag{F20}$$

The form of $K(t)$ is that of a simple exponential kernel with a delay (Fig. 4B). We thus have both $K(0) = 0$ and $K(\infty) = 0$. It is now clear that the convolution over the ligand concentration simply maps onto the convolution over its derivative as

$$\int_{-\infty}^{0} k(-t)\ell(t)dt = -\int_{-\infty}^{0} K(-t)v(t)dt. \tag{F21}$$

The static gain of $K(t)$ is $\bar{g}_{v\to x} = \int_0^\infty K(t)dt = q\gamma\tau_{\rm r}\tau_{\rm m} = \kappa N X_{\rm T}(1-p)f(1-f)\tau_{\rm m}$. The gain thus increases with the number of receptors per cluster, $N$, the number of readout molecules, $X_{\rm T}$, and notably, with the adaptation time $\tau_{\rm m}$. This static gain from signal derivative to network output is a useful quantity which we will use to describe the other statistics of the network below.

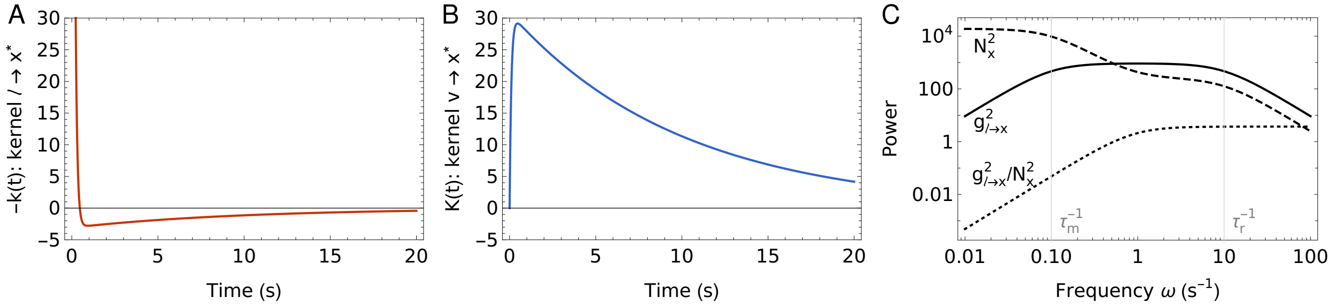


FIG. 4. **Integration kernel and power spectra.** (A) The integration kernel $k(t)$ takes a temporal derivative by weighing the most recent signal values with an opposite sign from the preceding ones. (B) The integration kernel $K(t)$ from the derivative of the input concentration to the network output. The kernel $K(t)$ is the primitive of $k(t)$, and its static gain is proportional to the adaptation timescale $\tau_{\rm m}$. (C) Frequency dependent gain $\tilde{g}^2_{\ell\to x}(\omega)$, frequency dependent noise $N_x^2(\omega)$, and their ratio, as a function of frequency. The chemotaxis network is a band-pass filter, the frequencies that are passed through are set by $\tau_{\rm r}$ on the high end and $\tau_{\rm m}$ on the low end. At low frequencies, the methylation noise dominates. Parameters used in all panels $\tau_{\rm r} = 0.1$s and $\tau_{\rm m} = 10$s. Model parameters are $\tilde{a} = 2$, $N = 6$, $K_{\rm D}^{\rm I} = 18\mu$M, $K_{\rm D}^{\rm A} = 2900\mu$M, $\bar{\ell} = 100\mu$M, $p = f = 0.5$.

To compute the past and predictive information, we need to determine the variance in the output, the SNR, and the correlation between the current output and the future ligand derivative. To that end we require the power spectrum of the output, and the cross-spectrum from output to future derivative. For the power spectrum of the output we use Eq. A5 to find

$$S_x(\omega) = \frac{q^2\gamma^2\omega^2}{(\tau_{\rm r}{}^{-2}+\omega^2)(\tau_{\rm m}{}^{-2}+\omega^2)} S_\ell(\omega) + \frac{\alpha^2\gamma^2\langle\eta_m^2\rangle}{(\tau_{\rm r}{}^{-2}+\omega^2)(\tau_{\rm m}{}^{-2}+\omega^2)} + \frac{\langle\eta_m^2\rangle}{\tau_{\rm r}{}^{-2}+\omega^2}. \tag{F22}$$

From this power spectrum we can see that the network is a band-pass filter, where the gain is maximal in the frequency range $\tau_{\rm m}{}^{-1} < \omega < \tau_{\rm r}{}^{-1}$. Both for $\omega \gg \tau_{\rm r}{}^{-1}$ and $\omega \ll \tau_{\rm m}{}^{-1}$ the gain goes to 0. On long timescales the methylation noise dominates (Fig. 4C). The cross-power spectrum between current output and future ligand derivative is given by element $(2,2)$ of the matrix $\mathbb{G}(-\omega)\mathcal{S}_s(\omega)$ which is (also see Eq. A6)

$$S_{x\to v}(\omega) = q\gamma \frac{-\omega^2 S_\ell(\omega)}{(\tau_{\rm m}{}^{-1} - i\omega)(\tau_{\rm r}{}^{-1} - i\omega)}. \tag{F23}$$

In the main text, we argue that the biologically relevant regime of the input signal is the limit $\omega_0 \to 0$. We therefore present below the network statistics in this limit. We start by determining the variance in the readout, via the inverse Fourier transform of its power spectrum (Eq. F22):

$$\begin{aligned}\lim_{\omega_0\to 0}\sigma_x^2 &= \bar{g}^2_{v\to x}\frac{1+\tau_{\rm r}/\tau_v+\tau_{\rm r}/\tau_{\rm m}}{(1+\tau_{\rm m}/\tau_v)(1+\tau_{\rm r}/\tau_v)(1+\tau_{\rm r}/\tau_{\rm m})}\sigma_v^2 + \bar{g}^2_{a\to x}\alpha R_{\rm T} p(1-p)\frac{1}{1+\tau_{\rm r}/\tau_{\rm m}} + X_{\rm T} f(1-f),\\ &= \underbrace{\bar{g}^2_{v\to x}\frac{1+\tau_{\rm r}/\tau_v+\tau_{\rm r}/\tau_{\rm m}}{(1+\tau_{\rm m}/\tau_v)(1+\tau_{\rm r}/\tau_v)(1+\tau_{\rm r}/\tau_{\rm m})}}_{\text{dynamical gain}}\sigma_v^2 + X_{\rm T} f(1-f)\left(1+\bar{g}_{v\to x}\frac{\tilde{\alpha}(1-p)}{R_{\rm T}\kappa\tau_{\rm m}}\frac{1}{1+\tau_{\rm r}/\tau_{\rm m}}\right),\end{aligned} \tag{F24}$$

where $\bar{g}_{a\to x} = \gamma\tau_{\rm r} = X_{\rm T} f(1-f)/(R_{\rm T}p)$ is the static gain from receptor activity to readout, and we used the definition of $\alpha = \tilde{\alpha}Np(1-p)$. Because there is no receptor-ligand binding noise, there is also no time averaging as in the push-pull network (and hence no factor depending on $\tau_{\rm r}/\tau_{\rm c}$). There is methylation noise on a timescale $\tau_{\rm m}$, but this cannot be time-averaged effectively because the integration time $\tau_{\rm r}$ of the push-pull network is shorter than the receptor methylation timescale $\tau_{\rm m}$. The methylation noise can only be averaged out significantly by increasing $R_{\rm T}$. The contribution from the variance in the signal derivative, $\sigma_v^2$, to the output noise $\sigma_x^2$, depends on the dynamical gain, which is the product of the static gain $\bar{g}^2_{v\to v}$ and a factor that only depends on ratios of timescales. The dynamical gain is maximized for $\tau_{\rm r} \to 0$ and $\tau_{\rm m} \to \infty$, which is intuitive since subtracting a signal from an earlier one reduces the amplification of the signal. Hence, when the system has too few $X_{\rm T}$ molecules to lift the signal above the noise, $\tau_{\rm m}$ must be increased to raise the gain. Only when $X_{\rm T}$ is sufficiently large, can $\tau_{\rm m}$ be reduced. This allows the system to take more recent derivatives. The signal to noise ratio SNR $= \sigma^2_{x|\eta}/\sigma^2_{x|L}$ can straightforwardly be obtained from Eq. F24. For the covariance between the current output and the future derivative we have

$$\lim_{\omega_0\to 0}\langle \delta x(0)\delta v(\tau)\rangle = \mathcal{F}^{-1}\{S_{x\to v}(\omega)\},$$
$$= \frac{-\bar{g}_{v\to x}\sigma_v^2}{(1+\tau_{\rm m}/\tau_v)(1+\tau_{\rm r}/\tau_v)} e^{-\tau/\tau_v}. \tag{F25}$$

The variance in Eq. F24 can be used to obtain the normalized correlation function $\langle \delta x(0)\delta v(\tau)\rangle/(\sigma_x\sigma_v)$.

#### 4. Past and predictive information of the chemotaxis network

The past and predictive information are straightforward to compute from the quantities above. The definition of the past information is the same as for the push-pull network, and is given by Eq. D3. The SNR is now given by, using Eq. F24:

$$\text{SNR} = \sigma^2_{x|\eta}/\sigma^2_{x|L} = \kappa^2 N\tau_m^2\sigma_v^2 \frac{1+\tau_{\rm r}/\tau_v+\tau_{\rm r}/\tau_{\rm m}}{(1+\tau_{\rm m}/\tau_v)(1+\tau_{\rm r}/\tau_v)(1+\tau_{\rm r}/\tau_{\rm m})} \Bigg/ \left(\frac{1}{NX_{\rm T}f(1-f)(1-p)^2} + \frac{\tilde{\alpha}}{R_{\rm T}(1+\tau_{\rm r}/\tau_{\rm m})}\right),$$

where $\kappa = (\bar{\ell}+K_{\rm D}^{\rm I})^{-1} - (\bar{\ell}+K_{\rm D}^{\rm A})^{-1}$. The predictive information is found in the same manner as in Eq. D6, but now it is a function of the correlation between the current output and the future *derivative* of the ligand concentration. This correlation can be decomposed into the instantaneous correlation coefficient and an exponential decay on the timescale of the fluctuations of the derivative of the concentration, Eq. F25. Specifically, the predictive information is given by

$$I_{\rm pred}(x_0; v_\tau) = -\frac{1}{2}\log(1-\rho^2_{\ell v}e^{-2\tau/\tau_v}). \tag{F26}$$

The instantaneous correlation coefficient $\rho^2_{\ell v}$ can be found using Eq. F25 and Eq. F24. From Eq. F26 it is clear that just like for the push-pull network, the optimal design of the network that maximizes the predictive information, determined by the optimal ratio $X_{\rm T}/R_{\rm T}$ and the optimal adaptation time $\tau_{\rm m}$, does not depend on the forecast interval $\tau$. The forecast interval only affects the magnitude of the predictive information.

#### 5. Optimal allocation

We can determine the optimal ratio $(X_{\rm T}/R_{\rm T})^{\rm opt}$ that maximizes either the past information or the predictive information, given all other network parameters, most notably $\tau_{\rm m}$. Just as for the push-pull network, we find however that the optimal ratio $(X_{\rm T}/R_{\rm T})^{\rm opt}$ is the same regardless of whether the past or the predictive information is maximized. This is again because the information on the future signal (be it the value or the derivative) is encoded in the receptor occupancy, while the ratio $X_{\rm T}/R_{\rm T}$ controls the interval by which the downstream readout samples the receptor to estimates its occupancy. Nonetheless, the optimal methylation timescale ${\tau_{\rm m}}^{\rm opt}$ that maximizes either the past or the predictive information is different—maximizing predictive information requires a more recent derivative and hence a shorter $\tau_{\rm m}$ than obtaining past information.

Given $\tau_{\rm m}$ and all other parameters, the optimal ratio of the number of readout molecules over receptor clusters is, using $C = R_\mathrm{T} + X_\mathrm{T}$,

$$\left(\frac{X_\mathrm{T}}{R_\mathrm{T}}\right)^\mathrm{opt} = \sqrt{\frac{1}{\alpha}\frac{1}{f(1-f)}\frac{p}{1-p}}\sqrt{1+\frac{\tau_\mathrm{r}}{\tau_\mathrm{m}}}, \\ = 2\sqrt{2/N}\sqrt{1+\frac{\tau_\mathrm{r}}{\tau_\mathrm{m}}}, \tag{F27}$$

where in the second line we have used that $\alpha = \tilde{\alpha}Np(1-p)$, and $\tilde{\alpha} = 2$, and $f = p = 0.5$. Because for the chemotaxis network $\tau_\mathrm{r} < \tau_\mathrm{m}$ the ratio $\tau_\mathrm{r}/\tau_\mathrm{m}$ only varies between 0 and 1. For this reason, the optimal ratio $(X_\mathrm{T}/R_\mathrm{T})^\mathrm{opt}$ depends only weakly on $\tau_\mathrm{m}$, and does not vary strongly along the isocost lines of Fig. 4A in the main text, see Fig. 5.

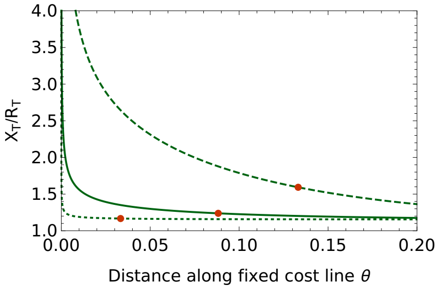


FIG. 5. **The optimal allocation ratio $X_\mathrm{T}/R_\mathrm{T}$ varies only slightly along the isocost lines of Fig. 4A in the main text.** The optimal ratio $X_\mathrm{T}/R_\mathrm{T}$ as a function of the distance $\theta$ along the isocost lines of Fig. 4A of the main text; dotted line $C = 10^2$, solid line $C = 10^4$, dashed line $C = 10^6$. The red dots mark the points where the predictive information is maximal. Along the isocost lines $X_\mathrm{T}/R_\mathrm{T}$ varies much more weakly than for the push-pull network; for resource availability $C \leq 10^4$ the ratio is almost constant. Parameters used $g = 4\mathrm{mm}^{-1}$, $\tau_\mathrm{r} = 0.1s$, $K_\mathrm{D}^\mathrm{I} = 18\mu\mathrm{M}$, $K_\mathrm{D}^\mathrm{A} = 2900\mu\mathrm{M}$, $N = 6$, $\tilde{\alpha} = 2$, $p = f = 0.5$, $\bar{\ell} = 100\mu\mathrm{M}$.


[1] N. Van Kampen, *Stochastic processes in physics and chemistry* (North Holland, Amsterdam, 1992).
[2] S. Tanase-Nicola, P. B. Warren, and P. R. ten Wolde, Signal detection, modularity, and the correlation between extrinsic and intrinsic noise in biochemical networks, Phys. Rev. Lett. **97**, 068102 (2006).
[3] E. Ziv, I. Nemenman, and C. H. Wiggins, Optimal signal processing in small stochastic biochemical networks., PLoS ONE **2**, e1077 (2007).
[4] W. De Ronde, F. Tostevin, and P. R. ten Wolde, Effect of feedback on the fidelity of information transmission of time-varying signals, Phys. Rev. E **82**, 031914 (2010).
[5] M. Vennettilli, S. Saha, U. Roy, and A. Mugler, Precision of Protein Thermometry, Physical Review Letters **127**, 098102 (2021), publisher: American Physical Society.
[6] H. H. Mattingly, K. Kamino, B. B. Machta, and T. Emonet, Escherichia coli chemotaxis is information limited, Nature Physics 2021 17:12 **17**, 1426 (2021), publisher: Nature Publishing Group.
[7] V. Sourjik and H. C. Berg, Functional interactions between receptors in bacterial chemotaxis, Nature 2004 428:6981 **428**, 437 (2004), publisher: Nature Publishing Group.
[8] B. A. Mello and Y. Tu, Effects of adaptation in maintaining high sensitivity over a wide range of backgrounds for Escherichia coli chemotaxis, Biophysical Journal **92**, 2329 (2007), publisher: Biophysical Society.
[9] N. Tishby, F. C. Pereira, and W. Bialek, The information bottleneck method, Proceedings of 37th Allerton Conference on communication and computation (1999).
[10] G. Chechik, A. Globerson, N. Tishby, and Y. Weiss, Information Bottleneck for Gaussian Variables, Journal of Machine Learning Research **6**, 165 (2005).
[11] C. C. Govern and P. R. ten Wolde, Optimal resource allocation in cellular sensing systems, Proceedings of the National Academy of Sciences **111**, 17486 LP (2014).
[12] G. Malaguti and P. R. T. Wolde, Theory for the optimal detection of time-varying signals in cellular sensing systems, eLife **10**, 1 (2021), publisher: eLife Sciences Publications Ltd.
[13] J. R. Maddock and L. Shapiro, Polar Location of the Chemoreceptor Complex in the Escherichia coli Cell, Science **259**, 1717 (1993), publisher: American Association for the Advancement of Science.
[14] T. A. J. Duke and D. Bray, Heightened sensitivity of a lattice of membrane receptors, Proceedings of the National Academy of Sciences **96**, 10104 (1999).
[15] T. S. Shimizu, Y. Tu, and H. C. Berg, A modular gradient-sensing network for chemotaxis in Escherichia coli revealed by responses to time-varying stimuli, Molecular Systems Biology **6**, 1 (2010), publisher: Nature Publishing Group.
